# Arabidopsis GENOMES UNCOUPLED PROTEIN1 binds to plastid RNAs and promotes their maturation

**DOI:** 10.1101/2024.02.08.579428

**Authors:** Qian Tang, Duorong Xu, Benjamin Lenzen, Andreas Brachmann, Madhura M Yapa, Paymon Doroodian, Christian Schmitz-Linneweber, Tatsuru Masuda, Zhihua Hua, Dario Leister, Tatjana Kleine

## Abstract

Plastid biogenesis and the coordination of plastid and nuclear genome expression through anterograde and retrograde signaling are essential for plant development. GENOMES UNCOUPLED1 (GUN1) plays a central role in retrograde signaling during early plant development. The putative function of GUN1 has been extensively studied, but its molecular function remains controversial. Here, we evaluate published transcriptome data and generate our own data from *gun1* mutants grown under signaling relevant conditions to show that editing and splicing are not relevant for GUN1-dependent retrograde signaling. Our study of the plastid (post)-transcriptome of *gun1* seedlings with white and pale cotyledons demonstrates that GUN1 deficiency significantly alters the entire plastid transcriptome. By combining this result with a PPR code-based prediction and experimental validation by RNA immunoprecipitation experiments, several targets of GUN1 were identified, including 23S rRNA, tRNAs and RNAs derived from *ycf1.2* and the *ndhH*-*ndhA*-*ndhI*-*ndhG*-*ndhE*-*psaC*-*ndhD* gene cluster. The absence of plastid rRNAs and the significant reduction of almost all plastid transcripts in white *gun1* mutants account for the cotyledon phenotype. Our study identifies RNA binding and maturation as the long-sought molecular function of GUN1 and resolves long-standing controversies. We anticipate that our findings will serve as a basis for subsequent studies investigating the mechanism of plastid gene expression and will facilitate the elucidation of GUN1’s function in retrograde signaling.

## Introduction

Chloroplasts are the characteristic organelles of algae and plants, and it is generally accepted that they are derived from ancient cyanobacteria through endosymbiosis (1). During evolution, most of the genes of the endosymbiont were transferred to the nuclear genome, resulting in only about 100 genes present in current plastid genomes (2) and at least 3,000 plastid proteins that are encoded in the nucleus (3). As a result, most plastid multiprotein complexes, such as the plastid gene expression (PGE) machinery and the photosynthetic apparatus, are formed by a mixture of plastid- and nuclear-encoded proteins, requiring the co-ordination of the expression of both genomes. Since most plastid proteins are encoded in the nucleus, it exerts anterograde control over the plastids. For example, the process of PGE necessitates the involvement of diverse nuclear-encoded proteins that promote the transcription, splicing, trimming, and editing of RNA in organelles, while simultaneously regulating their translation (4–7). On the other hand, nuclear gene expression, such as the expression of the so-called photosynthesis-associated nuclear genes (PhANGs), is controlled by plastid-to-nucleus retrograde signaling (8, 9), which is thought to be mediated by multiple factors and sources. For instance, in seedlings treated with norflurazon (NF) or lincomycin (LIN), mRNA levels of *PhANGs* are repressed (10). NF is an inhibitor of carotenoid biosynthesis (10), while LIN targets the peptidyl transferase domain V of the 23S rRNA of the 50S ribosomal subunit, which is the site of peptide bond formation, thereby preventing peptide bond formation (11). A mutant screen with *Arabidopsis thaliana* (Arabidopsis hereafter) identified a group of *genomes uncoupled* (*gun*) mutants three decades ago (12). In these mutants, the expression of *PhANG*s, in particular the marker gene *LHCB* encoding light-harvesting chlorophyll a/b-binding proteins of photosystem (PS) II, is de-repressed in seedlings treated with inhibitor (12). The original gun screens (12, 13) discovered six *gun* mutants, five of which, *gun2* to *gun6*, are impaired in the tetrapyrrole biosynthesis pathway. The *gun1* mutant exhibits a distinct *gun* phenotype when treated with LIN, distinguishing it from the other mutants (summarized in (14)). *GUN1* encodes a chloroplast pentatricopeptide repeat (PPR) protein (15). PPR proteins belong to a large family, with an estimated 106 of these proteins targeted to chloroplasts (6). They participate in various PGE steps, including RNA cleavage, splicing, editing, stabilization, and translation (6, 7). Thus far, no other *ppr* mutant has been identified as a *gun* mutant, indicating that GUN1 is a special component of an anterograde-retrograde axis.

GUN1 is an ancient protein that evolved within the streptophyte clade of the algal ancestors of land plants before the first plants colonized land more than 470 million years ago. It has been suggested that the primary role of GUN1 is to act in PGE, and that its involvement in retrograde signaling probably evolved more recently (16). In fact, GUN1 contains two domains known to interact with nucleic acids, namely the PPR domain and a MutS-related (SMR) domain (15). Among the large number of PPR proteins, Arabidopsis contains only eight PPR-SMR proteins, five of which are predicted to be localized in chloroplasts (17), including PLASTID TRANSCRIPTIONALLY ACTIVE 2 (pTAC2), SUPPRESSOR OF VARIEGATION 7 (SVR7), EMBRYO DEFECTIVE 2217 (EMB2217), SUPPRESSOR OF THYLAKOID FORMATION 1 (SOT1), and GUN1. Mutants of the first four show severe molecular and/or visible phenotypes, but only SOT1 has been shown to have an RNA-binding function (17, 18). Mainly by studying either *gun1* seedlings grown on inhibitors or in combination with other mutants, GUN1 has been implicated in a variety of processes in chloroplasts, to name a few, such as regulation of tetrapyrrole biosynthesis (19), protein homeostasis (20), ribosome maturation (21), accumulation of certain chloroplast transcripts and chloroplast import (22). Recently, it has been proposed that GUN1 cooperates with MULTIPLE ORGANELLAR RNA EDITING FACTOR 2 (MORF2)/ DIFFERENTIATION AND GREENING-LIKE 1 (DAL1) to regulate RNA editing under NF conditions (23). In the suggested mechanism GUN1 would not directly bind to the target RNAs. Rather, it would facilitate differential editing through its interaction with MORF2. Although GUN1 has been suggested to interact with DNA in vitro (15), no function in nucleic acid binding has yet been demonstrated in vivo. Furthermore, apart from occasional observations of pale cotyledons in a proportion of seedlings (e.g. in (24)), no clear severe phenotype has been observed.

In this study, we revisit the editing functions of GUN1 and MORF2 during retrograde signaling, define a distinct *gun1* phenotype with white cotyledons but green true leaves, examine the *gun1* (post)transcriptome in detail, and demonstrate an RNA binding function of GUN1.

## Results

### GUN1 does not play a significant role in plastid RNA editing or splicing during retrograde signaling

Based on the analysis of Sanger sequencing data, it has been proposed that GUN1 regulates plastid RNA editing during retrograde signaling (23). Previously, RNA sequencing after ribosomal RNA (rRNA) depletion (lncRNA-Seq) data covering both nuclear and organellar transcripts, were generated for wild-type (WT) and *gun1-102* seedlings grown on MS and NF (25). The benefit of the lncRNA-Seq technique is that its workflow involves library preparation after depletion of rRNAs instead of enriching for mRNAs, the latter approach applied by (23) and many other studies analyzing *gun1* mutants. Analysis of sequences generated by (25) for splicing and editing changes revealed no significant alterations between WT and *gun1-102* when grown on MS (**SI Appendix, Fig. S1 *A* and *B***). NF had a significant (secondary) impact on plastid splicing, and was similarly reduced in WT and *gun1-102* (**SI Appendix, Fig. S1*C***). Also, no major differences in editing (C-to-U base substitutions) efficiencies were observed between *gun1-102* and WT grown on MS (**SI Appendix, Fig. S1*D***), consistent with previous findings (23). Editing was reduced at multiple sites in NF-treated WT (**Fig. 1*A***), confirming that editing is altered under stress exposure (23, 26). According to Zhao et al. (23), GUN1-mediated editing is particularly important under inhibitor treatment. They found that RNA editing levels in *gun1-8* and *gun1-9* increased for *clpP-559*, *ndhB-467*/*-836*, *ndhD-878*, and *rps12-i-58*, but decreased for *rpoC1-488*, *ndhF-290*, *psbZ-50*, and *rpoB-338/-551/-2432* in comparison to WT when grown on NF. We confirmed increasing editing levels in *gun1-102* for the same sites (**Fig. 1*A***), but observed only a moderate reduction in RNA editing at two sites, *psbZ-50* (87% in WT, 82% in *gun1-102*) and *rpoB-338* (87% in WT, 79% in *gun1-102*). To account for different growth and analysis conditions, we repeated the experiment in two different laboratories, using the growth conditions employed by (23). Laboratory 1 employed *gun1-102* in Sanger sequencing experiments (**Fig. 1*B***), while laboratory 2 included both *gun1-1* and *gun1-102* in amplicon sequencing experiments (**Fig. 1*C***). These experiments revealed no reproducible differences in editing efficiency between WT and *gun1* under NF conditions, except for a slight reduction in *rpoC1-488* and *rpoB-551* editing.

**Figure 1.**
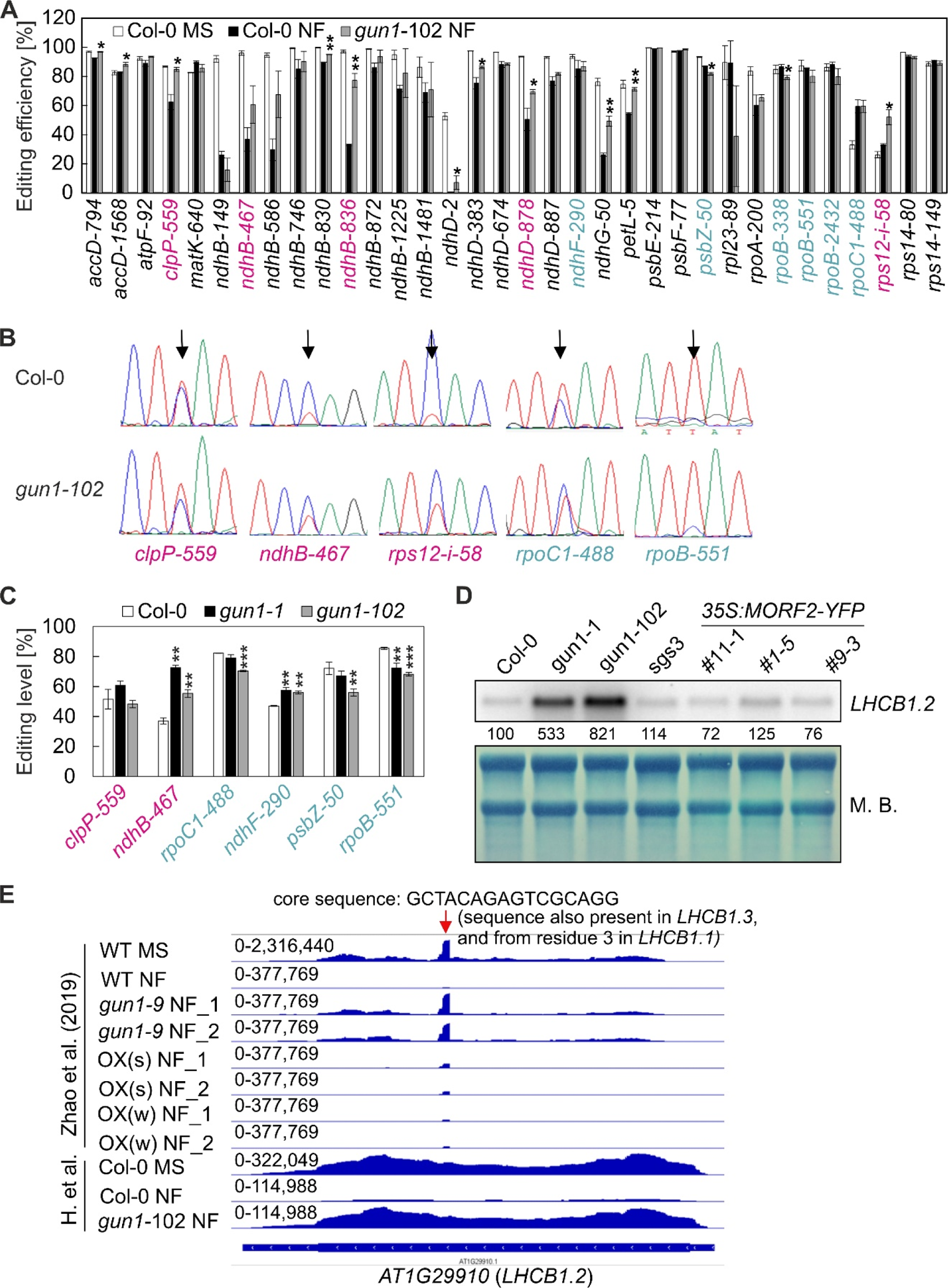
GUN1 does not play a significant role in plastid RNA editing or splicing during retrograde signaling. **(A)** RNA editing efficiencies of 4-day-old Col-0 and *gun1-102* seedlings grown on MS and norflurazon (NF) were determined using previously published RNA-Seq data (25). These sequencing data were generated to allow for detection of organellar transcripts. Mean values ± standard deviations were obtained from three independent experiments. Statistically significant differences between Col-0 NF and *gun1-102* NF are indicated (two-tailed student’s *t*-test; \**P* < 0.05, \*\**P* < 0.01). The efficiency of editing sites labeled in magenta and turquoise was found to be elevated and reduced, respectively, by Zhao et al. (23). We also identified an unexpected increase in editing of *rpoC1* in both WT and *gun1-102* under NF treatment. Our results may vary due to the utilization of different analysis methods - Sanger sequencing versus lncRNA-Seq data analysis, as well as discrepancies in growth media and conditions. Notably, Zhao et al. (23) cultivated five-day-old seedlings on MS plates without sucrose, while Habermann et al. (25) utilized MS plates with 1.5% sucrose. Thus, to account for these variations, we repeated the experiment for selected editing sites in two distinct laboratories as shown in panels B and C. **(B)** Col-0 and *gun1-102* seedlings were grown in Laboratory 1 for 5 days under continuous light conditions as reported by Zhao et al. (23). Editing efficiency of selected sites was visualized by Sanger sequencing. **(C)** Col-0, *gun1-1* and *gun1-102* seedlings were grown in Laboratory 2 for 5 days under continuous light conditions as reported by (23). Editing efficiency of selected sites was determined by amplicon sequencing. Shown are mean values with their standard deviations. Statistically significant differences between Col-0 and *gun1* seedlings are indicated (two-tailed student’s *t*-test; \**P* < 0.05 \*\**P* < 0.01; \*\*\**P* < 0.001). **(D)** Overexpression of MORF2 does not result in a significant *gun* phenotype. Shown are steady-state levels of *LHCB1.2* transcripts in 5-day-old seedlings grown under norflurazon (NF) conditions. Col-0 serves as the WT control for *gun1*, and *sgs3-1* as a control for oeMORF2 (*35S:MORF2-YFP*) lines. For each genotype, total RNA was fractionated on a formaldehyde-containing denaturing gel, transferred onto a nylon membrane, and probed with [α-32P]dCTP-labeled complementary DNA (cDNA) fragments specific for the transcripts encoding *LHCB1.2*. rRNA was visualized by staining the membrane with Methylene Blue (M. B.) and served as a loading control. Quantification of signals relative to the wild type (=100) is provided below each lane. **(E)** Snapshots of re-analyzed RNA-Seq data published by (23), and (25); H. et al.). The read depths were visualized with the Integrated Genome Browser (IGB). While reads from (25) are evenly distributed across *LHCB1.2*, reads generated by (23) exhibit a prominent peak of 16 nucleotides. The sequence of the peak (GCTACAGAGTCGCAGG) is also present in *LHCB1.3*, and from the third nucleotide, in *LHCB1.1*. The sequence of this peak coincidences with the sequence of the “LHB1.2” forward primer (actually detecting *LHCB1.3* in combination with the given reverse primer) used by (23) for qRT-PCR.

To summarize, the only mild editing and splicing differences between WT and *gun1* upon NF treatment argue against a major impact of these processes in GUN1 signaling.

### Overexpression of MORF2 does not result in a significant *gun* phenotype

Previously, two MORF2 overexpression lines *MORF2OX(s)* and *MORF2OX(w) MORF2OX(s)*, were constructed (23). Specifically, *MORF2OX(s)* exhibited a *gun* phenotype as its mRNA levels of nuclear-encoded photosynthesis genes, including *LIGHT HARVESTING CHLOROPHYLL A/B BINDING PROTEIN1.2* (*LHCB1.2*), were higher than those of WT when seedlings were treated with NF (23). We found that plants overexpressing *MORF2* in the Col-0 background induced co-suppression of *MORF2* and lead to variegation phenotypes in both early seedlings and adult plants (27), similar as observed for *MORF2OX(s)* (23). To prevent potential post-transcriptional co-suppression-mediated gene silencing, we introduced a *35S:MORF2-YFP* construct into *suppressor of gene silencing 3-1* (*sgs3-1*) plants (28) (**SI Appendix, Fig. S1**). At the cotyledon stage, lines *35S-MORF2-YFP #1-5* and *#11-1* exhibited phenotypes similar to that of Col-0 and *sgs3-1*. However, line *#9-3*, which had the highest induction of *MORF2* levels (**SI Appendix, Fig. S2*A***), displayed a reduction in the maximum quantum yield of photosystem II (Fv/Fm) (**SI Appendix, Fig. S2*B***). The determination of editing levels for *ndhF-290*, *psbZ-50*, *rpoB-338* and *rpoB-551*, sites that have been described to be less edited in both *MORF2OX(s)* and *gun1–9* seedlings under NF treatment (23), indicates that, interestingly, also in our strongest MORF2 overexpressor (*#9-3*) (**SI Appendix, Fig. S3*A***), editing levels for *ndhF-290* and *psbZ-50* are compromised compared to its parent plant *sgs3-1* (**SI Appendix, Fig. S3*B***).

To examine the *gun* phenotype of *35S:MORF2-YFP* lines, RT-qPCR was performed on retrograde marker genes. As expected, mRNA levels of the marker genes *LHCB1.2*, *CARBONIC ANHYDRASE 1* (*CA1*), and *PLASTOCYANIN* (*PC*) were higher in the *gun1* alleles. Although *LHCB1.2*, *CA1*, and *PC* mRNA levels were slightly elevated in line *#9-3*, they remained significantly lower than in *gun1* mutants and at a similar level to *sgs3-1* (**SI Appendix, Fig. S3*C***). Also, northern blot analysis showed high levels of *LHCB1.2* in *gun1* alleles, but WT-like levels in the *35S:MORF2-YFP* lines (**Fig. 1*D***). We reanalyzed the RNA-Seq data generated for WT, *gun1-9*, *oeMORF2(s)* and *oeMORF2(w)* (23) and the sequencing data from (25), and plotted the reads across the *LHCB1.2* gene. While the data from (25) showed even distribution of reads across *LHCB1.2*, reads generated by (23) exhibited a prominent peak of 16 nucleotides (**Fig. 1*E***). It is predominantly this peak that is found in MORF2 overexpressors after NF treatment (23), while there are almost no reads for the remainder of the *LHCB1.2* gene.

Overall, evidence suggests that overexpression of MORF2 does not result in a significant *gun* phenotype.

### The nuclear transcriptome of white and marble *gun1* seedlings is significantly affected

During the experiments examining the role of GUN1 in NF-mediated editing changes, we observed the appearance of *gun1* seedlings with white (*gun1W*) and marble (*gun1M*) cotyledons among the green *gun1* (*gun1G*) seedlings, when grown on MS medium without supplementation of inhibitors (**Fig. 2*A***), which was also previously reported (24), but to lower frequencies, which we will discuss later. The phenotype was most pronounced in *gun1-102* seedlings, but was also observed in *gun1-1* and *gun1-103* seedlings. The emerging true leaves turned green, suggesting that GUN1 especially has a role in chloroplast development in cotyledons which is in accordance with the particular accumulation of GUN1 protein levels at early stages of cotyledon development (29). To get a general overview of the RNA expression patterns in these prominent *gun1* seedlings, RNAs isolated from four-day-old Col-0 and *gun1W,* -*M*, and -*G* mutant seedlings (**Fig. 2*B***) were subjected to lncRNA-Seq. The absence of transcription in a portion of exon 2 and subsequent exons of the *GUN1* gene was verified in all *gun1* mutant seedlings (**SI Appendix, Fig. S4**), confirming the T-DNA insertion in all *gun1* seedlings and validating the RNA-Seq data. The strong phenotype of *gun1W* seedlings in particular suggests that the post(transcriptome) may be pleiotropically affected. Both NF and lincomycin (LIN) treated seedlings are bleached to the same degree as *gun1W* seedlings. Therefore, the following analyses involve data previously generated from NF-treated (25) and LIN-treated (30) seedlings to account for putative pleiotropic effects in *gun1W* seedlings. Furthermore, the reads from these published data sets were analyzed using the same methodology as for our own data. The severity of the *gun1* phenotype correlated with an increased number of deregulated genes (**Fig. 2*C***). In *gun1W* seedlings, the expression of 3349 genes (including chimeras) changed significantly compared to Col-0 (>2-fold, *P* < 0.05; **Dataset S1**). Of these, 1637 were decreased and 1712 were increased, while the number of de-regulated genes in *gun1M* and *gun1G* were 3188 and 830, respectively (**Fig. 2 *C* and *D***). The number of significant changes in gene expression in Col-0 following LIN or NF treatment was 3348 and 2100, respectively, which fell within the range or were lower than those observed in *gun1W* (**Fig. 2*E***). The degree of overlap between the gene sets of *gun1W*, LIN, and NF was significant, yet surprisingly low (**Fig. 2*F***) even though the phenotypes are similar. When examining the mRNA expression of the marker gene *LHCB1.2*, it showed only a mild decrease compared to the significant reduction in Col-0 seedlings treated with LIN or NF. This pattern was evident for nearly all of the *LHC* members (**Fig. 2*G***).

**Figure 2.**
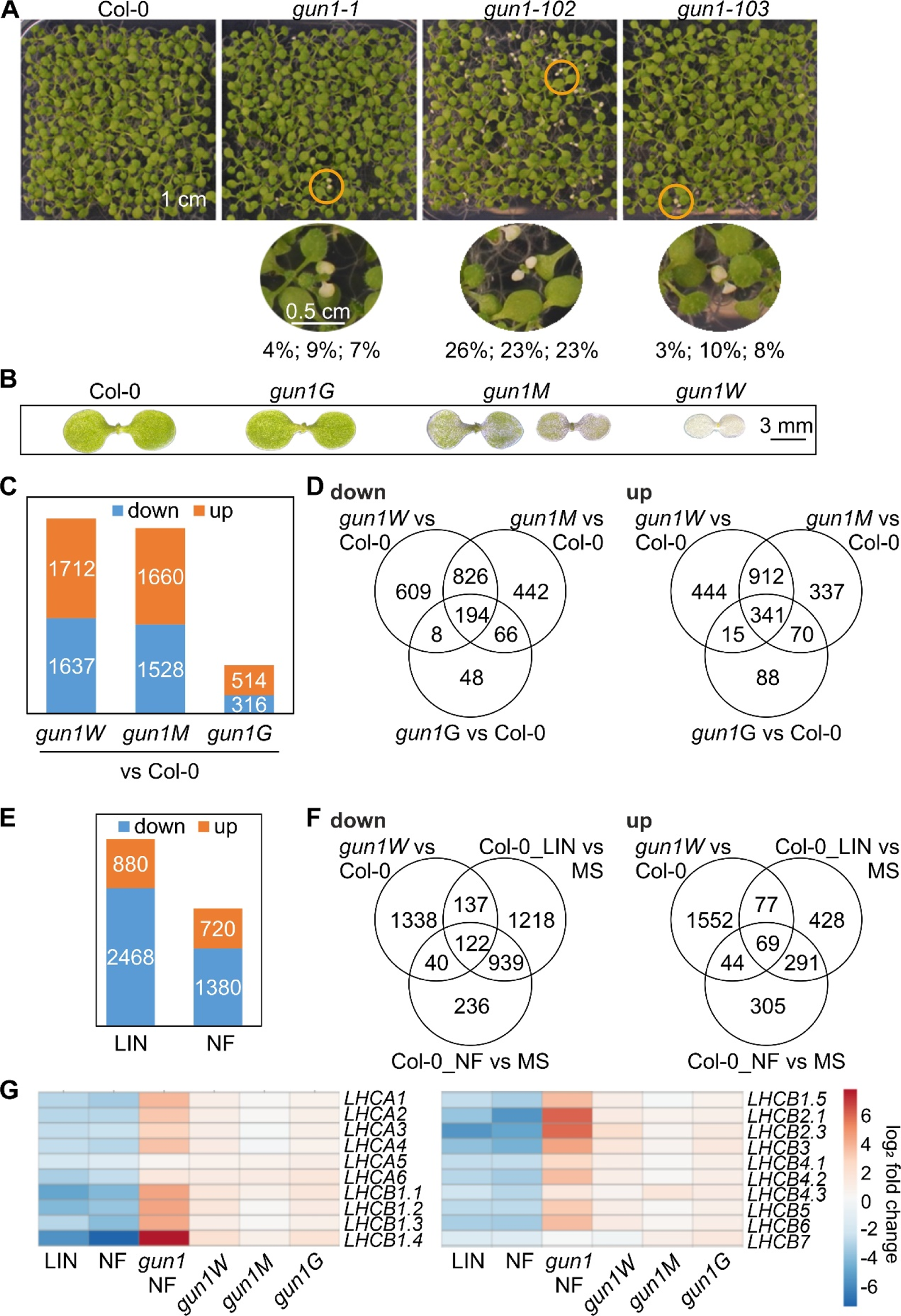
The nuclear transcriptome of white and marble *gun1* seedlings is significantly affected. **(A)** Phenotypes of 10-day-old Col-0, *gun1-1*, *gun1-102*, and *gun1-103* seedlings grown on MS without inhibitor supplementation under LD (16 h light, 8 h dark) conditions. Zoom-in images were taken of white seedlings, denoted by the circles below the overview pictures. The percentages of white seedlings were calculated for three different seed batches. **(B)** Phenotypes of Col-0, *gun1G*, *gun1M*, and *gun1W* seedlings (derived from *gun1-102*). **(C)** Analysis of transcriptome changes in white (*gun1W*), marble (*gun1M*), and green (*gun1G*) *gun1-102* mutant seedlings. The numbers represent the genes with at least a two-fold reduction (down), or elevation (up), in comparison with the Col-0 wild-type control. **(D)** Venn diagrams depicting the degree of overlap between the sets of genes whose expression levels were altered by at least two-fold in *gun1W*, *gun1M*, and *gun1G* as compared to the Col-0 control. **(E)** Re-analysis of transcriptome changes in lincomycin (LIN) (30) and norflurazon (NF) (25) treated seedlings. The numbers represent the genes with at least a two-fold reduction (down), or elevation (up), in comparison with the Col-0 control which was grown on medium without inhibitor. **(F)** Venn diagrams depicting the degree of overlap between the sets of genes whose expression levels were altered by at least two-fold in *gun1W* relative to Col-0, as well as in LIN and NF-treated seedlings as compared to Col-0 grown on medium without inhibitor (MS). **(G)** Heatmap showing transcript accumulation of genes encoding chlorophyll *a*/*b* binding proteins.

To summarize, the lack of GUN1 in *gun1W* and *gun1M* seedlings has a substantial effect on the nuclear transcriptome, but *LHC* transcripts are only mildly decreased.

### GUN1 deficiency has a significant impact on the entire chloroplast transcriptome

For the plastid transcriptome, we aimed to identify loci in which the relative ratio of editing or splicing was lower in *gun1W*/Col-0, progressively rescued in *gun1M*/Col-0 and *gun1G*/Col-0, and absent in NF/MS or LIN/MS. We concluded that the absence of GUN1 does not result in significant changes in chloroplast splicing or editing events (**SI Appendix, Figs. S5** and **S6**). However, plastid transcript levels of 68 out of 99 transcripts (including chimera) were significantly reduced in *gun1W* compared to WT, while no transcript was significantly induced (**Dataset S2**). Transcription of chloroplast genes relies on plastid-encoded (PEP) and nuclear-encoded polymerase (NEP) (5, 31). No clear conclusion can be drawn about PEP- or NEP-dependent transcription in *gun1W*: the so-called PEP-dependent genes were lower in *gun1W* compared to Col-0, as were the genes transcribed by PEP and NEP, although to a lesser extent. NEP-dependent gene expression was also reduced or in the range of Col-0 (**SI Appendix, Fig. S7**, **Table S2**). When we examined the down-regulated transcripts of *gun1W*, NF-, and LIN-treated Col-0 seedlings in parallel, we observed that remarkably, 21 transcripts were exclusively decreased in *gun1W* (**Fig. 3*A*; SI Appendix, Fig. S8*A***), four transcripts in NF and LIN data did not meet the threshold criteria and were not present in the gene sets (**Dataset S3**), but coverage plots showed that these transcripts were also reduced by LIN (**SI Appendix, Fig. S8*B***). Apart from few genes, most plastid genes belong to polycistronic units and are co-transcribed (32). A closer look at transcripts exclusively reduced in *gun1W*, gradually increased in *gun1M*, and WT-like in *gun1G*, drew our attention to a large polycistron containing *rpoA* along with several *rps* and *rpl* genes (**SI Appendix, Figure S8*C***). Inspection of coverage plots and transcript accumulation data revealed a comparable behavior for the *ycf1.2*, *rps15* and the *ndhH*-*ndhA*-*ndhI*-*ndhG*-*ndhE*-*psaC*-*ndhD* gene cluster (**Fig. 3*B***). In addition, during the quality control of RNA for sequencing, we observed strong rRNA depletion in *gun1W*, which was gradually rescued in *gun1M* and completely restored in *gun1G* (**Fig. 3*C***, **SI Appendix, Fig. S9*A***). The rRNA depletion phenotype was similar to that of Col-0 seedlings treated with NF or LIN (**Fig. 3*C***). Therefore, a secondary effect cannot be excluded at this stage. In conclusion, the entire plastid (post)transcriptome is significantly affected by GUN1 deficiency in *gun1W* and *gun1M* seedlings.

**Figure 3.**
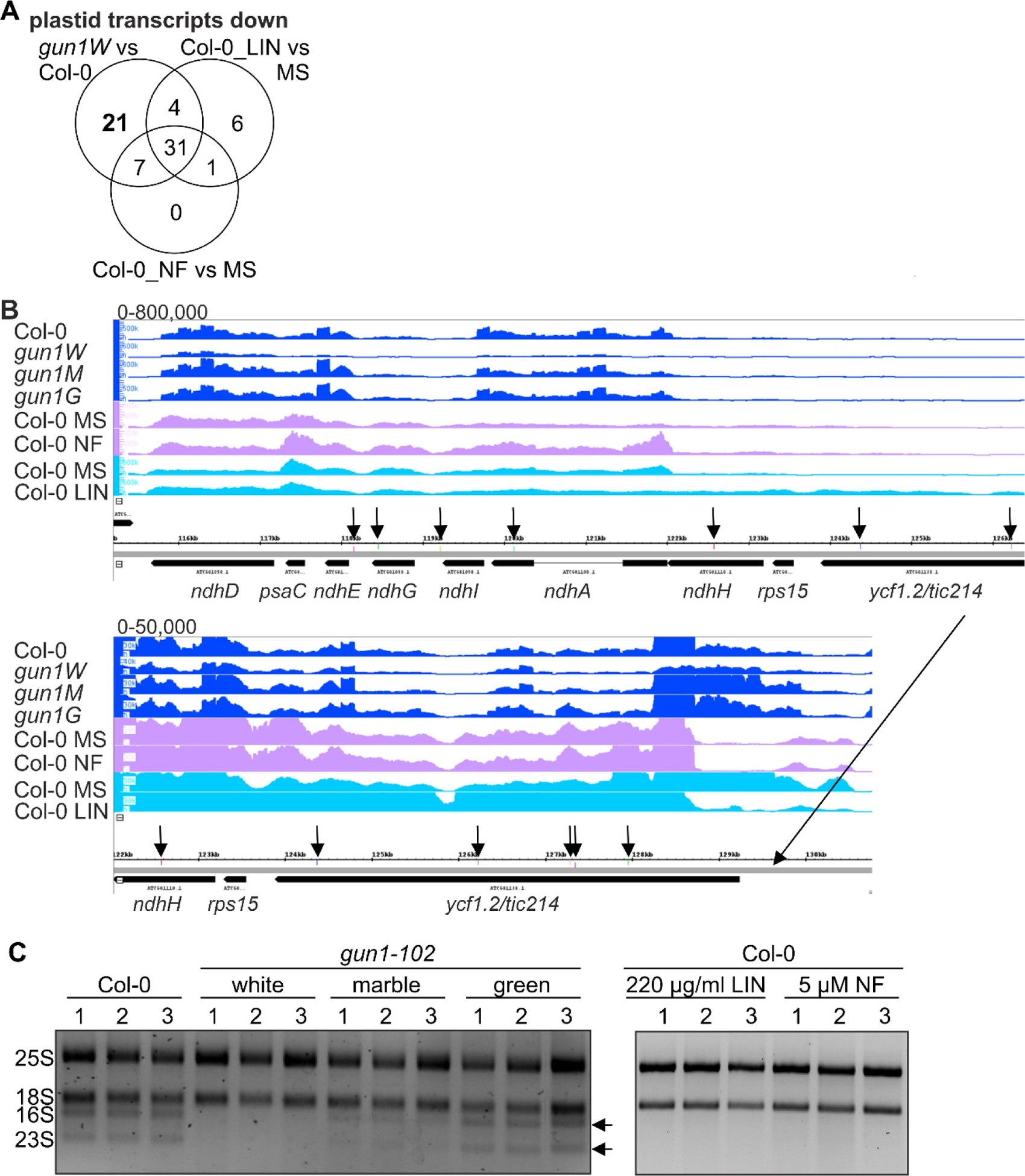
GUN1 deficiency has a significant impact on the entire chloroplast transcriptome. **(A)** Venn diagrams depicting the degree of overlap between the sets of plastid transcripts whose expression levels were reduced by at least two-fold in *gun1W* relative to Col-0, as well as in LIN and NF-treated seedlings as compared to Col-0 grown on medium without inhibitor (MS). **(B)** Coverage plots depict the accumulation of reads across the *ycf1.2*-*rps15*-*ndhH*-*ndhA*-*ndhI*-*ndhG*-*ndhE*-*psaC*-*ndhD* gene cluster. Vertical arrows point to predicted GUN1 binding sites (see Figure 6 and Supplemental Table 4). **(C)** Agarose gel displaying rRNA accumulation of RNAs isolated from 4-day-old Col-0, *gun1-102* white, marble and green seedlings, as well as from Col-0 seedlings grown on medium supplemented with norflurazon (NF) or lincomycin (LIN). The arrows point to the bands representing chloroplast rRNAs.

In conclusion, the entire plastid (post)transcriptome is significantly affected by GUN1 deficiency in *gun1W* and *gun1M* seedlings.

### Re-evaluation of a putative RNA binding function of GUN1

Many of the significant changes observed in the chloroplast (post)transcriptomes of *gun1W* and *gun1M* could explain their seedling phenotypes. But what is the primary cause? GUN1 is a P-type PPR protein, indicating that it may be associated with RNA cleavage, splicing, and stabilization (33), which led us to revisit a putative direct RNA binding function of GUN1. PPR motifs bind to RNA in a one-repeat, and one-nucleotide manner, and PPR motifs recognize specific RNA bases through amino acids at positions 5 and 35. Utilizing this code, the binding sites of several PPR proteins can be predicted very well (34–36). Since the correct PPR code is crucial for determining the binding sequence, we investigated the structural configuration of the GUN1 protein by modeling with PyMOL, and found that the 12 PPR domains of GUN1 predicted by ScanProsite should be shifted by one amino acid (**Fig. 4*A***), and adjusted the repeat annotation to better fit the predicted structure and description of canonical PPR tracts (16, 36). Predicting putative RNA target sites (36) yielded the following ambiguous 11-nucleotide sequence: 5’-AA(U>C>G)(U>C>G)(C>U)(G>>C)(U>C>G)(C>U)(G>>C)A(C>U>A)-3’ (**Fig. 4*B***). Using this sequence and considering location in inverted repeat regions, 78 potential targets can be identified within the chloroplast genome (**Dataset S4**). The application of strict and very strict sequence matching criteria yields 25 and 9 possible targets, respectively. Based on our previous analysis, two regions are noteworthy: the *ycf1.2*-*rps15*-*ndhH*-*ndhA*-*ndhI*-*ndhG*-*ndhE*-*psaC*-*ndhD* gene cluster (**see Fig. 3*B***), which contains ten potential targets. Among these targets, *ndhE* and 3’*ndhI* are also identified with the strict, and *ndhG* with the very strict target sequence, respectively (**Fig. 4*B***). The second region is the *rrn23S* gene (**Fig. 4 *B*** and ***C***), which comprises four predicted target sequences: 23S_104766, 23S_104856, 23S_106002, and 23S_106558 (numbering according to the nucleotide position in the cp genome). 23S_104856 and 23S_106558 fall within the strict possible targets. To gain insight into the accumulation of reads across the ribosomal RNA (rRNA) operon, we conducted lncRNA-Seq again without rRNA depletion. This confirmed that plastid rRNAs are significantly reduced in *gun1W* and *gun1M* seedlings (**SI Appendix, Fig. S9*B***). Upon closer examination of the first two binding sites and the adjustment of plots for differences in expression, a disproportionately high number of reads were found to map 5’ to the *rrn23S* gene, which is not present in *gun1G* (**Fig. 4*C***). Additionally, a distinct coverage pattern of *rrn23S* was observed in the region of binding site 23S_106558.

**Figure 4.**
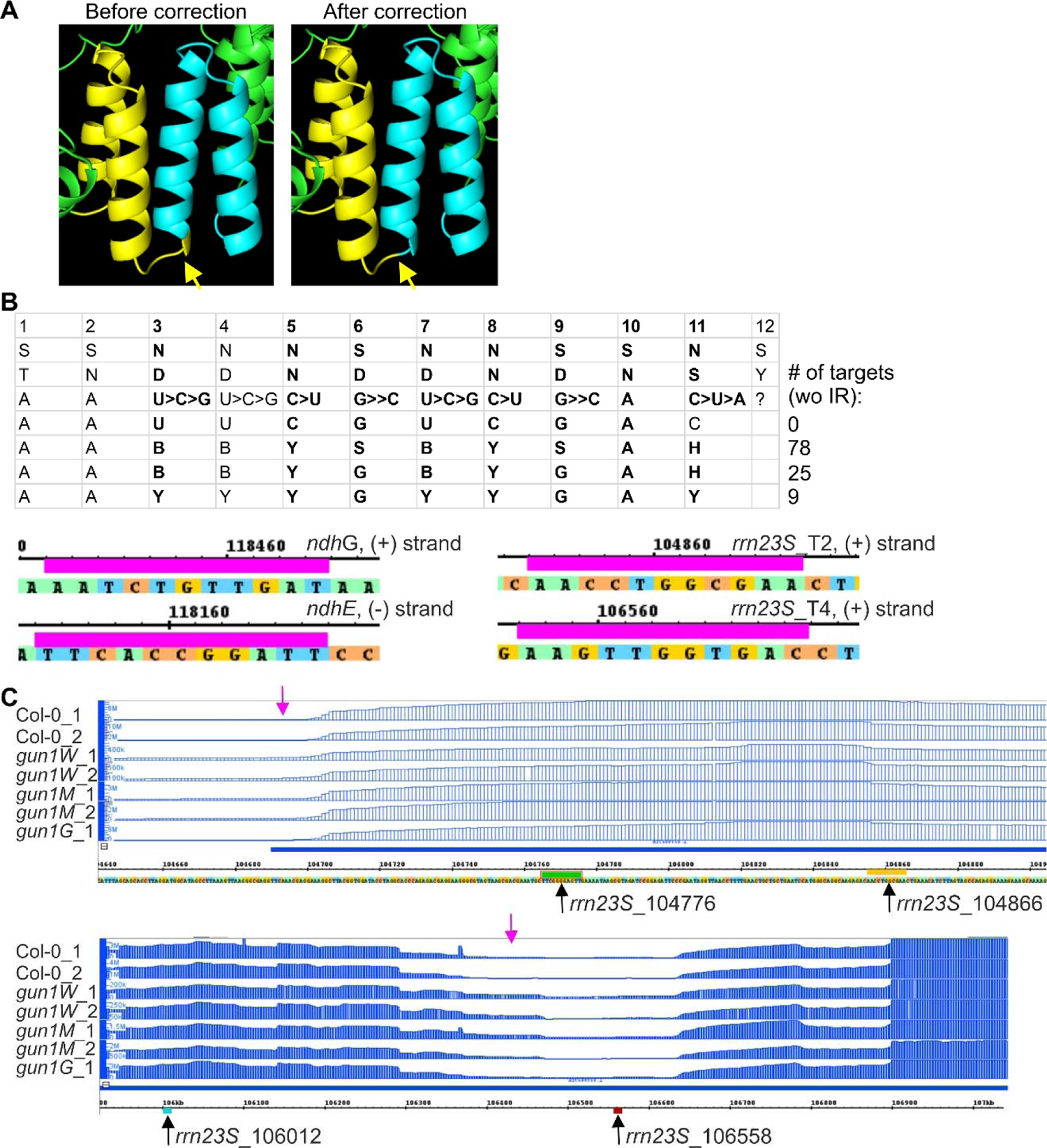
Predicted GUN1 binding sites and maturation of 23S ribosomal RNA. **(A)** Section of PPR domain modeling of the GUN1 protein. The 12 predicted PPR domains of GUN1 by ScanProsite and the PPR CODE PREDICTION WEB SERVER (http://yinlab.hzau.edu.cn/pprcode; (36)) differ by a shift of one amino acid. Since the correct PPR code is crucial for determining the binding sequence, we investigated the structural configuration of the GUN1 protein by modeling with PyMOL. Each PPR domain was marked with a unique color for easy differentiation and visualization. Notably, within this color-coded scheme, helix b (yellow) of the initial PPR domain is observed to extend beyond (marked with an arrow) helix a (turquoise) of the subsequent PPR domain. **(B)** Predicted ambiguous GUN1 target sequence. The numbers in the first row depict the PPR motif number, while the second and third rows display the amino acids in each PPR motif that are crucial in predicting target nucleotides. Subsequent rows indicate the prospective targets. The identification of possible targets in the chloroplast genome is dependent on the stringency of the utilized sequence, with 97, 31, or 10 being the potential targets. Highly conserved regions in GUN1 are highlighted in bold letters, according to (16). Additionally, representative predicted binding sites at *ndhG*, *ndhE*, and *rrn23S* are shown. wo IR, without inverted repeat. **(C)** Plots of RNA-Seq data produced without rRNA depletion depict a proportional increase in reads in *gun1W* and *gun1M* that map to 23S ribosomal RNA regions (magenta arrows) close to the four predicted GUN1 binding sites (marked with black arrows).

### GUN1 binds to chloroplast RNAs in vivo and in vitro

To investigate whether GUN1 is involved in RNA binding in vivo, RNA co-immunoprecipitation was carried out using a GPF-tagged GUN1 line (*GUN1-GFP*) (20). Col-0 and another unrelated GFP-tagged line (*PP7L-GFP*; (37)) served as controls. The success of the IP experiment was demonstrated by the detection of the tagged proteins in the respective eluates by Western blotting (**SI Appendix, Fig. S10**). Four predicted target regions of the notable regions described above (*ndhG*, *ycf1.2*, two regions of *23S* rRNA; **Fig. 5*A***) along with negative controls were tested in RT-qPCRs of immunoprecipitated RNAs. Both GFP-tagged lines were observed to bind more RNAs than Col-0, but this was not statistically significant for *PP7L-GFP*. In GUN1 IPs, *ndhG*, *ycf1.2*, and a target of *23S* rRNA comprising binding sites 104766 and 104856 demonstrated significant enrichment in the pellet compared to PP7L (**Fig. 5*B***). In contrast, there was no significant enrichment of RNAs without predicted target sites, except for *ycf3* with low fold enrichment. The qPCRs of the input RNAs did not show a significant enrichment for GUN1-GFP over Col-0 (**Fig. 5*B***). The binding of GUN1 to *23S*_106558 was less apparent and therefore unclear. Interestingly, the *ndhG* binding site of GUN1 falls within a region that appears to undergo intense maturation or degradation activity (38).

**Figure 5.**
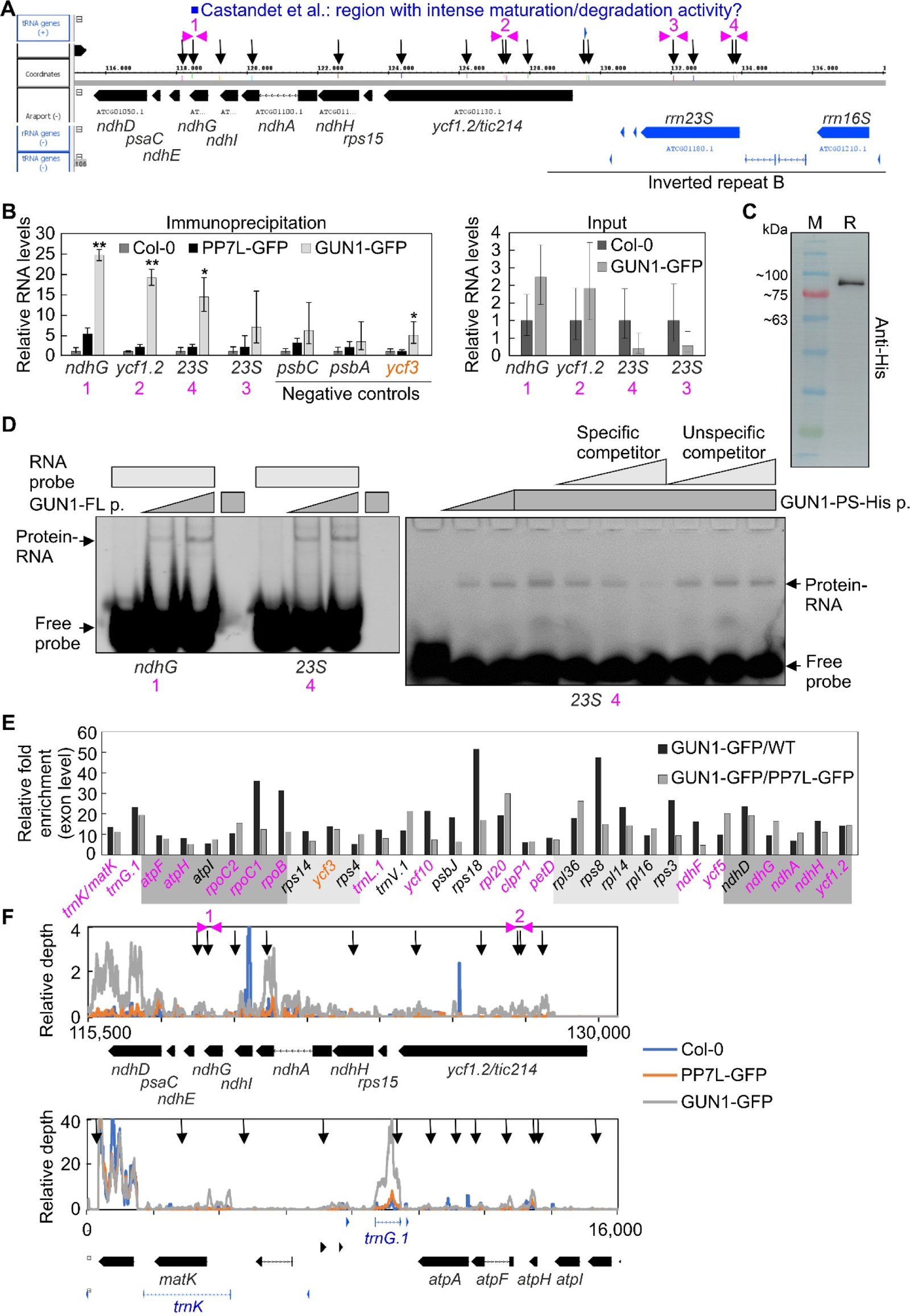
GUN1 binds to RNAs in vivo and in vitro. **(A)** Schematic presentation of predicted RNA binding sites (indicated by black vertical arrows) in *ycf1.2*, the *rps15*-*ndhH*-*ndhA*-*ndhI*-*ndhG*-*ndhE*-*psaC*-*ndhD* polycistron and the *rrn23S* gene. Positions of primers used in panel (B) are depicted by magenta arrowheads. The blue rectangle marks the region that was identified by (38) as a region of significant maturation or degradation activity. **(B)** Demonstration of co-purification of selected RNAs with GUN1. RNAs that were isolated from the pellet after co-immunoprecipitation experiments with Col-0, a PP7L overexpression line (PP7L-GFP), a GUN1 overexpression line (GUN1-GFP) (Immunoprecipitation), and the respective input RNAs (Input) were amplified by RT-qPCRs. RNA levels are reported relative to the corresponding levels in Col-0, which were set to 1. Mean values ± SD were derived from three independent experiments, each performed with three technical replicates per sample. Statistically significant differences (two-tailed student’s *t*-test; \**P* < 0.05; \*\**P* < 0.01) between GUN1-GFP and PP7L-GFP lines are indicated by black asterisks. **(C)** Western blot to demonstrate the purity of His-tagged GUN1 protein that was synthesized in vitro. **(D)** The GUN1 protein interacts in vitro with RNA sequences located in *ndhG* and *23S* rRNA. EMSAs were performed with purified His-tagged GUN1 proteins that were either synthesized in vitro (GUN1-FL) or produced in E. coli (GUN1-PS). Left: Aliquots (0, 200, 400 nM) of purified GUN1-FL protein (GUN1-FL p.) were incubated with Cy5-labelled ssRNA probes representing the target sequences. Right: Aliquots (0, 500, 1000, 1500 nM) of purified GUN1-PS protein (GUN1-PS-His p.) were incubated with Cy5-labelled ssRNA probes in the presence of increasing concentrations (indicated by the light gray triangle) of the same unlabeled ssRNA or an unlabeled ssRNA of unrelated sequence as competitors. Binding reactions were then subjected to electrophoresis on nondenaturing TBE-polyacrylamide gels. **(E)** Libraries were prepared from RNAs isolated from the co-immunoprecipitation experiments described in panel (B). Relative enrichment ratios (calculated at the exon level) of GUN1-GFP relative to Col-0 and GUN1-GFP relative to PP7L-GFP are shown. Gray shading indicates genes located in a polycistron. **(F)** Plot of RIP-Seq data over two example regions. Relative depth was calculated at each nucleotide (nt) position by relating the number of reads to the total depth of the sequencing output. Black vertical arrows indicate predicted GUN1 RNA binding sites.

To determine whether GUN1 can directly bind to the identified target sites, we used electrophoretic mobility shift assays (EMSAs). It is difficult to obtain full-length GUN1 overexpressed in *E. coli*, possibly due to the highly disordered domain in the N-terminal region (19). Therefore, we overexpressed the GUN1-PS construct encompassing all PPR and SMR motifs (PS) spanning amino acids 232 to 918 (19) in *E. coli* (**SI Appendix, Fig. S11**). In a second approach, we obtained the full-length GUN1 protein (GUN1-FL) by coupled vitro transcription/translation (**Fig. 5*C***). Both, GUN1-PS and GUN1-FL, were used for electrophoretic mobility shift assays (EMSAs). Two different Cy5-labeled oligonucleotides were designed, representing the binding sites *ndhG*, and *23S*_104856. When 200 and 400 nM of purified GUN1-FL protein were added to the Cy5-labeled probes and the mixtures electrophoresed, band shifts were observed for both probes. The shift was more pronounced at the higher protein concentration and was not detected when no protein or probe was added, indicating that the RNA probes formed complexes with the protein (**Fig. 5*D***). Furthermore, the intensity of the shifted *ndhG* and *23S*_104856 bands progressively decreased when increasing concentrations of the respective unlabeled ssRNAs were added to the incubation mixtures, employing GUN1-FL as well as GUN1-PS protein (**Fig. 5*C***; **SI Appendix, Fig. S12**). However, as shown for the *23S* binding site, the intensity of the shifted bands did not decrease when increasing concentrations of unlabeled unrelated ssRNA were added (**Fig. 5*D***), suggesting that GUN1 binds specifically to the target sites examined.

To get a broader view of the RNA targets of GUN1, libraries prepared from the immunoprecipitated RNAs were subjected to RNA sequencing. Enrichment analysis at the exon level compared to RNAs identified in the control lines identified 31 transcripts as significantly enriched (**Fig. 5*E***, **Dataset S5**); 18 of them contain predicted GUN1 target regions. These include *ycf1.2*, the *ndhH*-*ndhA*-*ndhI*-*ndhG*-*ndhE*-*psaC*-*ndhD* gene cluster and tRNAs such as *trnK* and *trnG.1* (**Fig. 5 *E*** and ***F***).

Overall, these experiments clearly demonstrate the RNA binding function of GUN1.

## Discussion

While the functions of other GUN proteins are well established, the specific molecular function of GUN1 has remained largely unclear. Most conclusions regarding GUN1 have been made by examining *gun1* mutants in combination with inhibitor treatments, or in conjunction with the generation of double mutants (14). Our observation of *gun1W* and *gun1M* seedlings is independent of NF or LIN treatment. Previous reports have also noted the sporadic presence of variegated or paler cotyledons (24, 29)). In *gun1W* and *gun1M* seedlings, the emerging true leaves turned green, suggesting a role of GUN1 especially in chloroplast development in cotyledons. This is consistent with the particular accumulation of GUN1 at early stages of cotyledon development (29).

### “Same genotype, different phenotype” phenomenon

The prevailing view concerning *gun1* mutants is that adult plants exhibit no noteworthy phenotypes under normal growth conditions, apart from earlier flowering (29, 39). Most inferences related to GUN1 were made when the *gun1* mutant was examined under stressful conditions, in combination with inhibitor treatments, or in conjunction with the creation of double mutants (see Introduction). We observed the appearance of *gun1* seedlings with white (*gun1W*) and marble (*gun1M*) cotyledons, when grown under normal growth conditions and without supplementation of inhibitors (see Fig. 3). Previous reports also already noted the sporadic presence of variegated (observed in *gun1-1* and *gun1-101*; (24)), or paler (observed in *gun1-101*; (29)) cotyledons. It is interesting to note that seedlings with the same genotype can exhibit varying phenotypes. This phenomenon, also described as incomplete penetrance and variable expressivity, is widely discussed in the animal field, also because of its relevance for diseases (40). Epigenetic modifications, and environmental effects are potential factors that could contribute to this occurrence. Environmental effects on mutants impaired in plastid gene expression have been observed, such as in the case of *gun1* mutants, which exhibit a defect in cold acclimation (39). However, we want to exclude a pure environmental effect for the appearance of *gun1W* seedlings since they are interspersed between green *gun1* seedlings on the same plate. Epigenetic changes, specifically DNA methylation and histone modifications, have the ability to impact gene expression without modifying the DNA sequence. These changes again can be influenced by environmental factors and can results in distinct phenotypes despite having identical genotypes. The *gun1W* seedlings were observed in diverse laboratories with different generations and different *gun1* alleles, including complete knock-outs (*gun1-101* and *gun1-102*). Therefore, also epigenetics may not be the primary/sole contributing factor. A comparable scenario to that of *gun1* mutants was described for the *immutants* (*im*) and *variegated2* (*var2*) mutants. Nevertheless, these mutants exhibit green and white sectors within the same leaf. The discussion of these mutants revolves around the compensatory mechanisms and the concept of plastid autonomy for both mutants. However, while redundant gene products are suggested to be involved in *var2*, they are not implicated in *im*. The hypothesis is that the attainment of certain activity threshold levels is imperative for the proper development of chloroplasts (41), which may also apply for the *gun1* mutants. A threshold effect would also explain the sensitivity of *gun1* mutants to LIN and NF (42, 43) and abscisic acid (44) during early seedling development. In addition, GUN1 protein accumulates at early stages of cotyledon development, and the timing of gene expression during development is known to influence penetrance (40). However, further analysis is needed and may include how stochastic factors – such as segregation of organelle genomes through development and reproduction (45) - in conjunction with environmental factors and transgenerational effects contribute to the development of individual phenotypes (46).

### Functions of GUN1 in plastid transcript maturation

GUN1 is suggested to regulate plastid RNA editing during NF treatment of seedlings (23). The proposed mechanism involves the interaction of GUN1 with MORF2 and does not require the direct interaction of GUN1 with the target transcript, which was a logical explanation because no in vivo RNA-binding function of GUN1 could be demonstrated so far. However, the role of GUN1 in editing and its contribution to GUN signaling have not yet been satisfactorily resolved for several reasons. First, the oeMORF2 *gun* phenotype has been postulated for NF treatment, but not for LIN treatment. Second, the slight differences in editing performance between Col-0 and *gun1* during NF treatment (see Fig. 1) are unlikely to be the trigger for retrograde signaling. Third editing of relevant sites was either more or equally suppressed in oeMORF2 compared to *gun1-9*, and consequently one would expect oeMORF2 lines to be even stronger *gun* mutants than *gun1* itself, which was not the case for both our data and data generated by (23) (see Fig. 1). Here it should be noted that our oeMORF2 lines had lower *MORF2* mRNA expression levels compared to those generated by (23). However, also other studies do not find any involvement of GUN1 in editing changes in other retrograde signaling processes (47, 48). Furthermore, GUN1 is classified as a P-type PPR protein, which rarely have a direct role in editing (49).

GUN1 is one of the five PPR-SMR cp-located proteins, and together with our study, it is demonstrated that all of them have essential functions in cp development (see Fig. 2; (17)). Interestingly, the GUN1 protein is present at very low levels and barely detectable by proteomic approaches, while the other PPR-SMR proteins are particularly abundant compared to most PPR proteins (50). This fact together with the distinct (post)-transcriptome in *gun1* mutants (see Figs. 3-5) may be important for the unique function of GUN1 in GUN signaling, as *svr7* and *sot1* mutants are no *gun* mutants (51). Interestingly, plastid rRNA accumulation is impaired in mutants of the three proteins SVR7, SOT1, and GUN1. While GUN1 (see Fig. 5) and SOT1 (18, 51) bind directly to the (precursor) *23S* rRNA, this is not clear for SVR7, and the defect in rRNA accumulation may be a secondary effect in the *svr7* mutant. However, the primary function of SVR7 is to ensure correct expression of the ATP synthase (52). For SOT1, specifically its function in rRNA maturation has been investigated, and it has been shown that the SMR domain has endonuclease activity (18, 51), but other targets are so far unknown. Interestingly, in contrast to the *gun1* mutant, the plastid transcripts of protein-coding genes (except *NdhA*) tend to be slightly up-regulated in *sot1* (36), whereas the *gun1* (post)-transcriptome is largely affected, and we have identified a plethora of enriched RNA sites in our RIP-Seq experiment (see Fig. 5).

We focused on the function of GUN1 in the maturation of *ycf1.2*/*tic214* and the *ndhH*-*ndhA*-*ndhI*-*ndhG*-*ndhE*-*psaC*-*ndhD* gene cluster. Interestingly, GUN1 is predicted to bind to multiple sites in *ycf1.2*, and no *ycf1.2* maturation factors have been identified to date. Furthermore, GUN1 binds to the *ndhG* region, which is assumed to undergo intense maturation or degradation activity (38). The significantly reduced *ycf1.2* (53) or plastid rRNA levels (54) would be sufficient to explain the *gun1W* phenotype. In addition, judging from RIP-Seq data, GUN1 could also be involved in tRNA maturation. Our data do not reveal precisely how GUN1 performs its function on plastid RNA, which may involve transcript stabilization or endonucleolytic cleavage through its SMR domain. In addition, we do not address how the molecular function of GUN1 relates to retrograde signaling. Nevertheless, we definitively established that GUN1 binds to RNA and identified its target sites. We anticipate that our findings will serve as a foundation for subsequent studies exploring the role of GUN1 in plastid RNA metabolism and retrograde signaling.

## Materials and Methods

### Plant material and growth conditions

The *gun1-1*, the T-DNA insertional mutants *gun1-102* (*SAIL_290_D09*) and *gun1-103* (*SAIL_742_A11*), are derived from the Col-0 ecotype and were published before (for example (19)).

To detect editing levels via RT-PCR, surface-sterilized seeds were sown on Murashige and Skoog (MS) plates containing 0.8% (m/v) agar. The seeds were then stratified for 4 days in the dark at 4°C. Seedlings were grown for 5 days at 22°C under continuous illumination (100 μmol photons m^−2^ sec^−1^) provided by white fluorescent lamps. For norflurazon treatment, MS medium was supplemented with or without 5 μM final concentration of norflurazon (Sigma-Aldrich, 34364).

For RNA sequencing and RIP experiments, surface-sterilized plant seeds were sowed on half-strength MS plates containing 1% sucrose. The plates were subsequently kept in the dark at 4°C for two days. Following stratification, the seedlings were grown under a photoperiod of 16 hours (light phase) and 8 hours (dark phase) at 22°C with a light intensity of 100 μmol photons m^−2^ sec^−1^ for four days.

### Generation of oeMORF2 transgenic lines

The *35S:MORF2-YFP* transgene was constructed into the pFGC5941 binary transformation vector as described previously (27). To avoid post-transcriptional co-suppression and to stabilize high expression of *MORF2-YFP*, *35S:MORF2-YFP* was transformed into a post-transcriptional gene silencing mutant, *suppressor of gene silencing 3-1* (*sgs3-1*) (55). Plants containing a single insertion of *35S:MORF2-YFP* were identified based on a 3:1 (resistant / sensitive) segregation ratio of T2 plants grown on 1/2 MS medium containing 15 mg/L phosphinothricin. Homozygous transgenic plants were obtained in T3 and further self-fertilized to generate T4 plants that were used for phenotypic analysis in this work.

### RNA preparation, cDNA synthesis and RT-qPCR

70 mg of plant material was frozen in liquid nitrogen and then crushed using a TissueLyser (Retsch, model: MM400). 1 ml of TRIZOL (Invitrogen, Carlsbad, CA, USA) and 200 µl of chloroform were used for RNA isolation according to the manufacturer’s instructions. Subsequently, RNA was precipitated from the aqueous phase using isopropyl alcohol, then the resulting RNA pellet was washed with 70% (v/v) ethanol, and finally dissolved in RNase-free water. After DNase I treatment (New England Biolabs, Ipswich, MA, USA), 10 µg of RNA were further cleaned up by RNA Clean & Concentrator-5 (Zymo Research, Irvine, USA; R1016). 500 ng of the purified RNA was employed to synthesize cDNA with the iScript cDNA Synthesis Kit (Bio-Rad). RT-qPCR analysis was performed on a Bio-Rad iQ5 real-time PCR instrument with the iQ SYBR Green Supermix (Bio-Rad). The primers used for this assay are listed in **SI Appendix**, **Table S6**.

### RNA editing analysis by amplicon sequencing

The same growth conditions as used by (23) were applied. Total RNA was isolated from agar plate-grown seedlings by acid guanidinium thiocyanate-phenol-chloroform-based extraction and purified from the aqueous phase using the Monarch RNA Clean Up Kit (New England Biolabs, Ipswich, MA, USA; NEB). Genomic DNA in the samples was removed using TURBO™ DNase (Thermo Fisher Scientific) followed by purification with the Monarch RNA Clean Up Kit (NEB). RNA (1 µg per sample) was transcribed to cDNA by the Protoscript®II reverse transcriptase (NEB). *clpP*, *psbZ*, *rpoC1*, *rpoB*, *ndhB*, *ndhF* amplicons were amplified with the Q5® polymerase (NEB) from all samples. Amplification specificity was assessed by agarose gel electrophoresis and amplicons were subsequently purified with the Monarch PCR & DNA Clean Up Kit (NEB). Resulting DNA concentrations were spectrophotometrically measured by NanoDrop. Equimolar amounts of all amplicons from a given sample were pooled and analyzed by the Amplicon-EZ service from Genewiz. The resulting 250 bp paired-end reads were mapped with the short-read aligner BBMap (https://sourceforge.net/projects/bbmap) to an amplicon specific reference. RNA editing was assessed from the mapped reads as described previously (56).

### RNA gel-blot analysis

Total RNA was isolated from five-day-old seedlings using TRIzol reagent (Thermo Fisher Scientific, Waltham, MA, USA). RNA samples were digested with DNase I (NEB) to remove genomic DNA. Then, five micrograms of total RNA were electrophoresed on a denaturing formaldehyde gel, transferred to a nylon membrane (Hybond-XL; GE Healthcare, Freiburg, Germany), and cross-linked by UV light. Hybridizations were performed at 65°C overnight according to standard protocols. Results were visualized using the Typhoon scanner (GE Healthcare, Freiburg, Germany).

### RNA editing and splicing analysis of lncRNA-Seq data

To ascertain the presence of edited and spliced transcripts from organelles from lncRNA-Seq datasets, the Chloro-Seq pipeline (57) was utilized with modifications described in (58).

### RNA sequencing (RNA-Seq) and data analysis

Total RNA from plants isolated with Trizol (Invitrogen, Carlsbad, USA) and purified with Direct-zol™ RNA MiniPrep Plus columns (Zymo Research, Irvine, USA) was subjected to sequencing as previously described (37). RNA-Seq reads were analyzed on the Galaxy platform (59) essentially as described (37), except that reads were first mapped with the gapped-read mapper RNA STAR (60) to generate the coverage plots in a subsequent step. Second, the reads were mapped with Salmon (61) to identify differentially expressed genes as described in (58), except that the updated AtRTD3-QUASI high-resolution transcriptome (62) was used as the reference transcriptome.

### Protein expression and electrophoretic mobility shift assays (EMSAs)

The coding sequence of GUN1 was amplified with primers listed in **SI Appendix, Table S6**, to introduce EcoRV and XhoI restriction sites, and cloned into pEU-E01-His-TEV. Plasmid DNA of pEU-E01-GUN1-His-TEV was isolated using the Qiagen Plasmid Midi Kit (#12143). A medium-volume cell-free protein expression reaction was then performed with 5 μg of the pEU-E01-GUN1-His-TEV plasmid using the WEPRO7240 Core Kit (ENDEXT® Technology) in a bilayer format according to the manufacturer’s instructions. Where indicated, synthesized GUN1-FL protein was purified using His-Tag Dynabeads™ (Invitrogen; 10103D) and confirmed by Western blotting using His-tag antibody (Sigma; SAB 1305538). The His-tagged GUN1-PS protein was produced essentially as described (19). We used 1 mM IPTG for induction and induced for 20 hours at 18°C. The protein was purified using Protino Ni-NTA agarose (Macherey-Nagel, #7450400-500).

For EMSA experiments, the indicated amounts of GUN1 protein were used as input for the binding reactions, which for GUN1-FL consisted of 4 µl of 5x binding buffer (50 mM Tris-HCl pH 7.5, 50 mM NaCl, 200 mM KCl, 5 mM MgCl2, 5 mM EDTA, 5 mM DTT, 0.25 mg/mL BSA, and 5% glycerol) and Cy5-labeled probe. For GUN1-PS, the binding buffer was modified by adding 2-mercaptoethanol to the binding buffer and replacing glycerol with Ficoll 400. For the competitor assay, increasing amounts of competitor were added to the binding reaction. After incubation for 25 minutes at 25°C, 1 µl of 50% glycerol (v/v) was added. The samples were then run on a 5% native polyacrylamide gel preconditioned for 1 hour at 80 V in 0.5X TBE (pH 8.3 for GUN1-FL, and pH 8.6 for GUN1-PS) containing 2.5% glycerol to remove any residual ammonium persulfate. Gel electrophoresis was performed at 60 V until adequate separation was achieved, followed by scanning for the Cy5 signal on a Typhoon FLA9500 scanner.

### RNA immunoprecipitation – sequencing and RT-qPCR

For RNA immunoprecipitation (RIP), we adapted a previously described method (63) with some modifications. Four-day-old seedlings grown on 1/2 MS medium were fixed with 1% formaldehyde for 20 min by vacuum infiltration. The fixation was stopped with 125 mM glycine for 5 minutes, again by vacuum infiltration. The seedlings were then washed 4 times by prechilled sterile ddH20, ground to a fine powder with liquid nitrogen, and stored at -80°C for later use. 250 mg of the ground plant sample was homogenized in 1 ml of RIP buffer. The composition of the RIP buffer was consistent with that of the original paper. Instead of preparing the beads-antibody conjugate, commercial GFP Trap Magnetic Agarose beads (gtma-20: ChromoTek) were utilized. 40 µl of GFP-Trap was initially washed three times with 400 µl of RIP buffer and then incubated with 800 µl of cleared lysate for 2 h. The remaining steps for immunoprecipitation, RNA release, and extraction were conducted following the procedure previously outlined (63). Western blot was performed for input, flow through and pull-down fraction of all samples with GFP polyclonal antibody (Invitrogen; A6455). DNA contamination was removed by 2U DNaseI (NEB; M0303S) then purified with the RNA clean and concentrator 5 kit (ZYMO; R1016).

For subsequent sequencing, the RNA was processed with the NEBNext Ultra II RNA Library Prep Kit from Illumina (NEB; E7770L). The libraries were then sequenced on an Illumina NextSeq1000 system and analyzed on the Galaxy platform (59). For RT-qPCR, 2 µl of purified RNA was reverse transcribed using the SuperScript IV Reverse Transcriptase Kit (Invitrogen; 18090050) with random hexamer priming. The cDNA synthesis reaction was performed under the following conditions: initial incubation at 23°C for 10 minutes, followed by reverse transcription at 55°C for 15 minutes for efficient cDNA synthesis. The reaction was then inactivated by heating at 80°C for 10 minutes. RT-PCR was performed on a Bio-Rad iQ5 real-time PCR instrument using SYBR Green Supermix (Bio-Rad; 1725274). All primer information is provided in **SI Appendix, Table S6**.

## Data availability

Sequencing data have been deposited in NCBI’s Gene Expression Omnibus (64) and are accessible under the GEO Series accession number GSE202931. Reads from experiments conducted by (25), (23), and (30) were retrieved from the NCBI Sequence Read Archive (numbers PRJNA557616 and PRJNA432917, respectively) and Gene Expression Omnibus (number GSE130337).

## Acknowledgments

We thank David Meinke for critical discussions, Michael Färberböck and Katrin Straßer for excellent technical assistance, Irma Racic for library preparation for RIP-Seq, and Helmut Blum and Stefan Krebs for sequencing the RIP libraries.

