## Supporting Information for "Arabidopsis GENOMES UNCOUPLED PROTEIN1 binds to plastid RNAs and promotes their maturation"

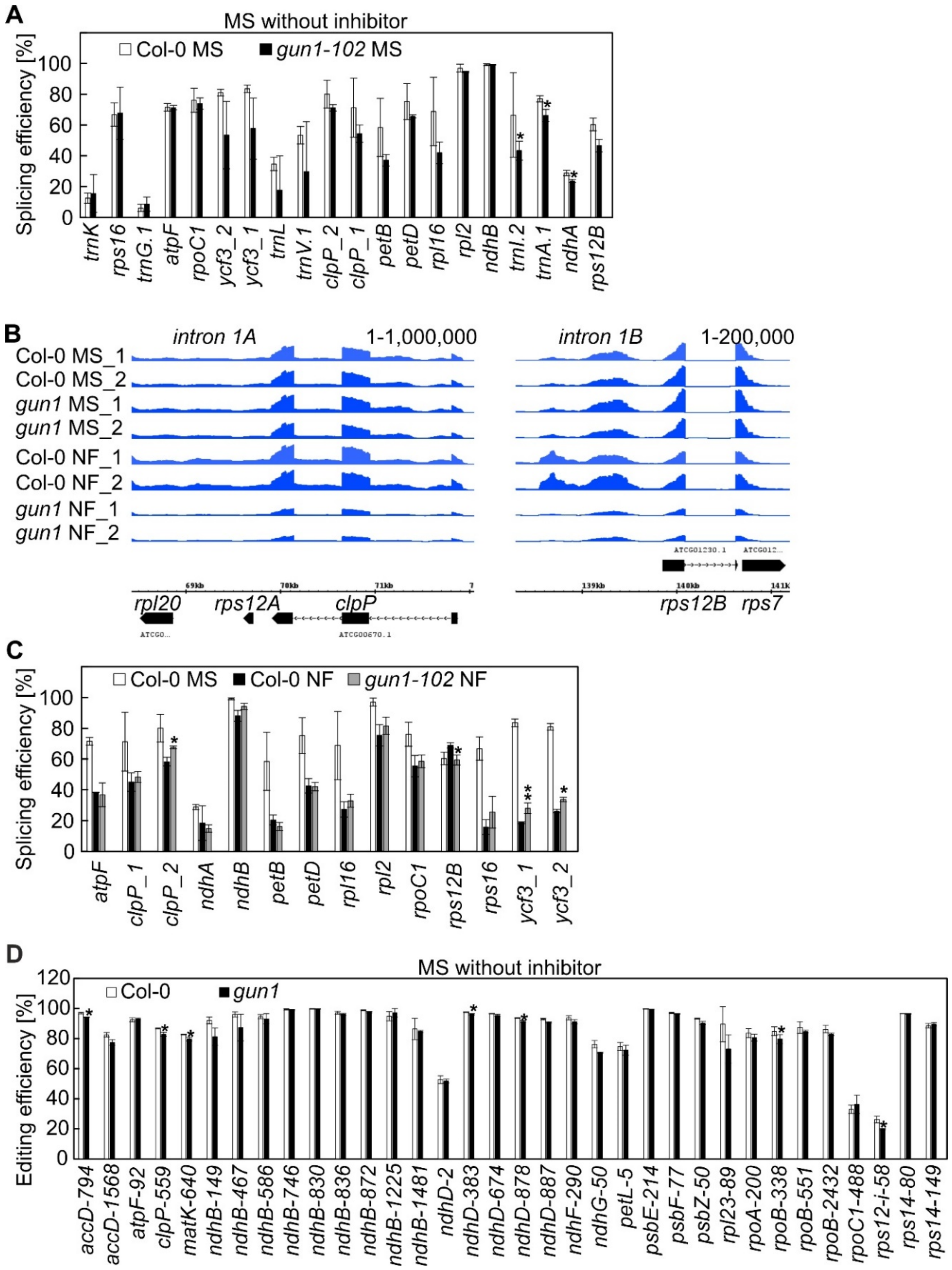


**Fig. S1. GUN1 does not play a significant role in plastid RNA editing or splicing under normal growth conditions.**

**(A)** RNA splicing efficiencies of 4-day-old Col-0 and *gun1-102* seedlings grown on MS were determined using previously published RNA-Seq data (1). These sequencing data were generated to allow for detection of organellar transcripts. Mean values ± SD were obtained from three independent experiments. Statistically significant differences between Col-0 MS and *gun1-102* MS are indicated (two-tailed student's *t*-test; *$P < 0.05$).

**(B)** Snapshots across the *clpP* gene and intron 1B of *rps12B*. The read depths were visualized with the Integrated Genome Browser (IGB). Intron 1 of *rps12* is transcribed from two separate chromosomal regions: one downstream of *rps12A* and the other upstream of *rps12B*. They are then spliced together in trans. Therefore, we conducted a manual investigation of this intron using coverage files from the sequencing data**.**

**(C)** RNA splicing efficiencies of 4-day-old Col-0 and *gun1-102* seedlings grown on MS and norflurazon (NF) were determined using previously published RNA-Seq data (1). These sequencing data were generated to allow for detection of organellar transcripts. Mean values ± SD were obtained from three independent experiments. Statistically significant differences between Col-0 NF and *gun1-102* NF are indicated (two-tailed student's *t*-test; *$P < 0.05$, **$P < 0.01$). Due to the numerous modifications, structure and small size of tRNAs, the lncRNA-Seq library preparation method is not reliable for their detection. Hence, the splicing efficiency data for the six tRNA introns have to be viewed with caution and were excluded from further analysis.

**(D)** RNA editing efficiencies were calculated from data described in (A).

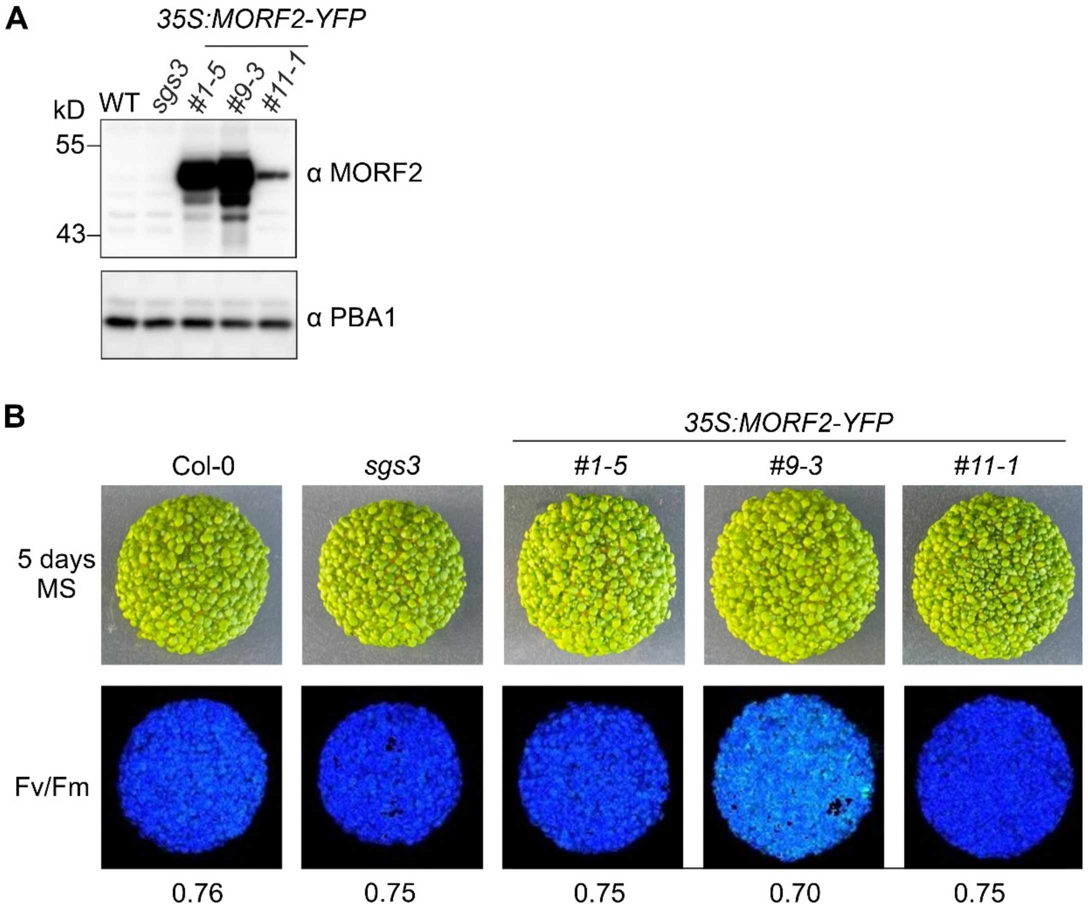


**Fig. S2. Characterization of *35S:MORF2-YFP* lines.**

**(A)** Immunoblot analysis showing protein levels of MORF2-YFP in three overexpression transgenic lines (*35S:MORF2-YFP*). Total proteins from 7-d-old seedlings were directly extracted in 2 x SDS sample buffer and denatured at 95°C for 6 min before being resolved on a 10% SDS-PAGE gel. MORF2-YFP was detected with an anti-MORF2 polyclonal antibody as described in (2). The 20S proteasome subunit PBA1 was used to verify nearly equal loading of total protein. The three lines, *35S-MORF2-YFP #1-5*, *#9-3* and *#11-1*, were selected for further analysis as they exhibit *MORF2* overexpression to varying degrees.

**(B)** Phenotypes and Fv/Fm Imaging PAM pictures of 5-day-old Col-0, *sgs3-1* and *35S:MORF2-YFP* lines grown on MS medium without inhibitor supplementation.

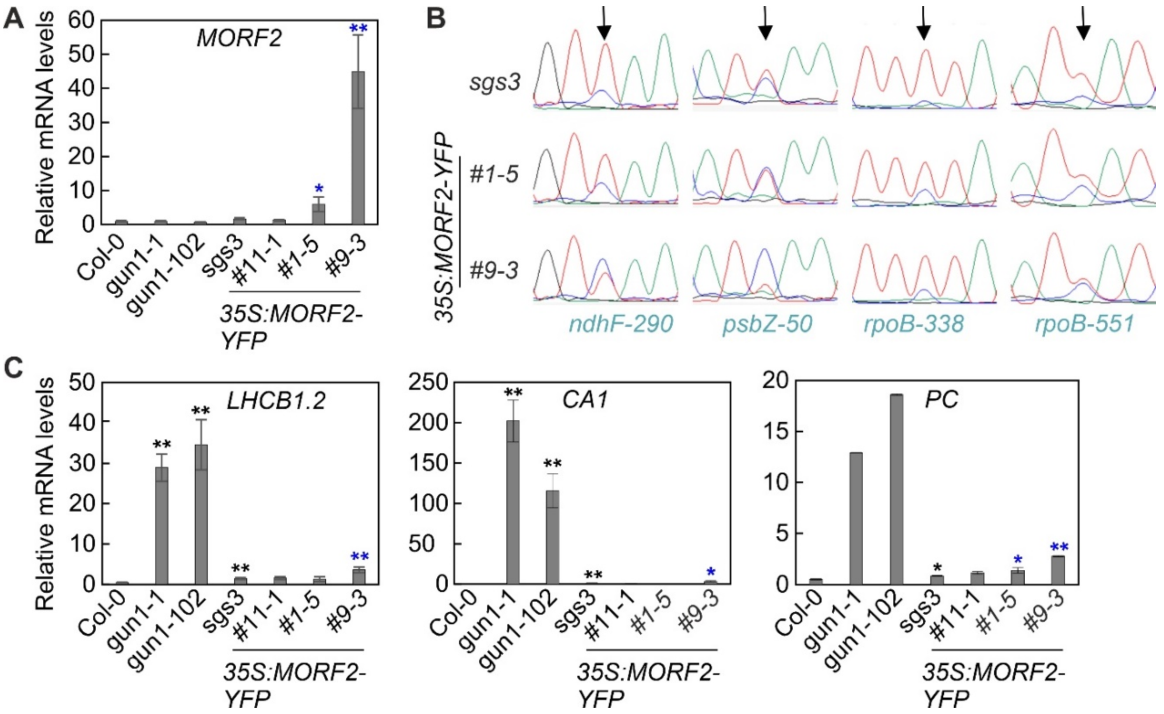


**Fig. S3. Overexpression of MORF2 does not result in a significant *gun* phenotype.**

**(A)** RT-qPCR of *MORF2* expression in 5-day-old seedlings grown under norflurazon (NF) conditions. The results were normalized to *AT4G36800*, which encodes a RUB1-conjugating enzyme (RCE1). Expression values are reported relative to the corresponding transcript levels in Col-0, which were set to 1. Mean values ± SE were derived from three independent experiments, each performed with three technical replicates per sample. Statistically significant differences (two-tailed student's *t*-test; **P* < 0.05; ***P* < 0.01) between Col-0, *gun1* and *sgs3-1* mutants are indicated by black asterisks, and those between *sgs3-1* and the *35S:MORF2-YFP* lines are indicated by blue asterisks.

**(B)** Seedlings were grown as in (A). Editing efficiency of selected sites was visualized by Sanger sequencing.

**(C)** RT-qPCR of *LHCB1.2*, *CARBONIC ANHYDRASE 1* (*CA1*), and *PLASTOCYANIN* (*PC*) was performed using the identical cDNAs as in (A).

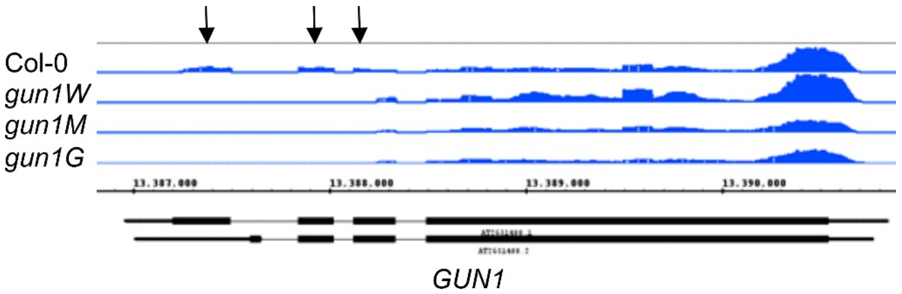


**Fig. S4. Validation of RNA-Seq data and the *gun1-102* allele.**
A snapshot across the *GUN1* gene is shown. The read depths were visualized with the Integrated Genome Browser (IGB). Arrows point to the absence of reads in a portion of exon 2 and the subsequent exons in *gun1W*, *gun1M*, and *gun1G*. The absence of transcription in a portion of exon 2 and subsequent exons of the *GUN1* gene was verified in all *gun1* mutant seedlings, confirming the T-DNA insertion in all *gun1* seedlings and validating the RNA-Seq data.

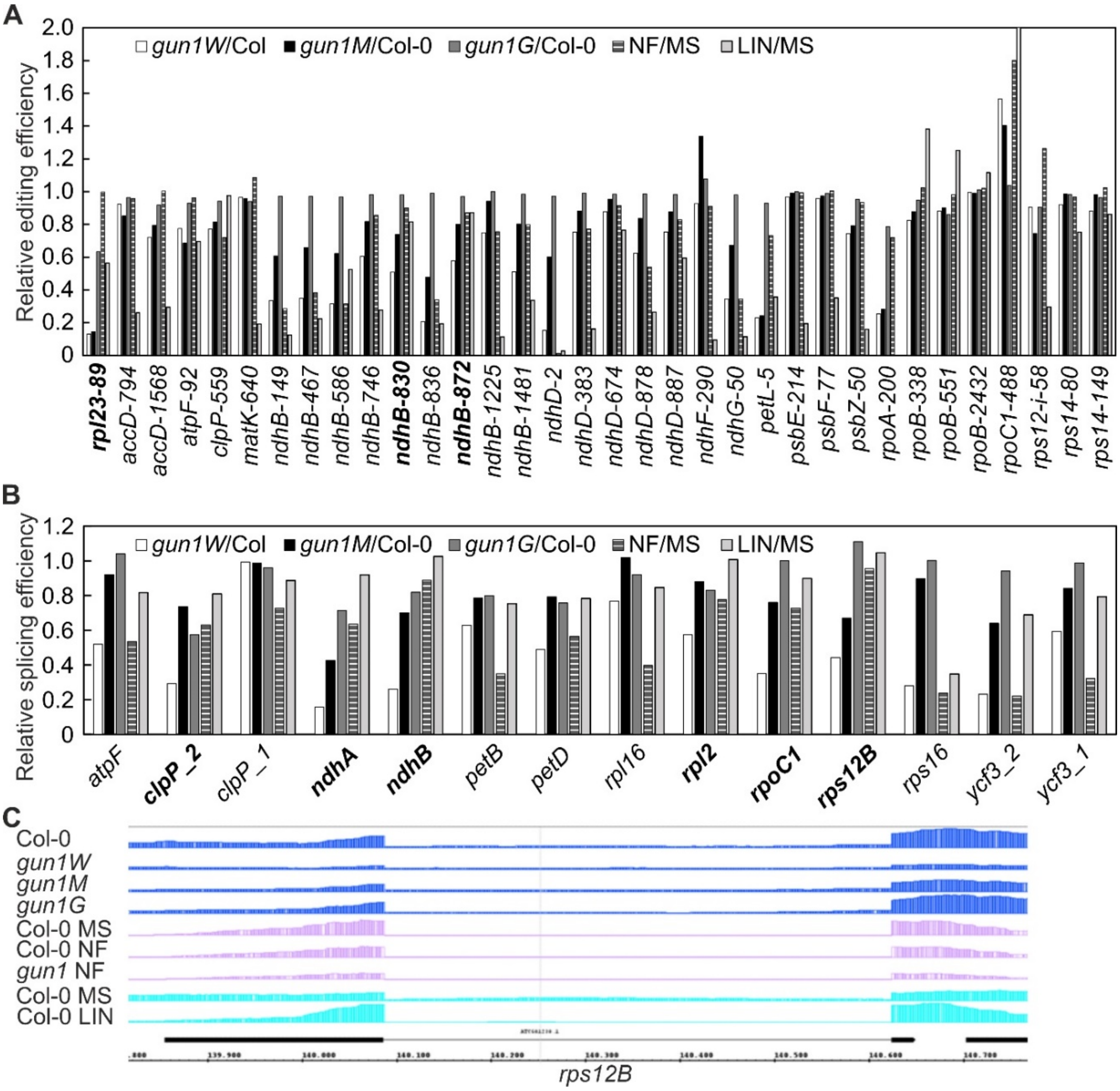


**Fig. S5. Analysis of editing and splicing efficiencies of *gun1W*, *gun1M* and *gun1G* seedlings in comparison to seedlings grown on norflurazon (NF) and lincomycin (LIN).**

**(A, B)** RNA editing **(A)** and splicing **(B)** ratios of 4-day-old white (*gun1W*), marble (*gun1M*), and green (*gun1G*) *gun1-102* seedlings compared to Col-0 (Col), and ratios of seedlings grown on norflurazon (NF) or lincomycin (LIN) compared to seedlings grown on MS. The NF, LIN, and MS data were extracted from previously published RNA-Seq data (1). We identified loci in which the relative ratio of editing or splicing was lower in *gun1W*/Col-0, progressively rescued in *gun1M*/Col-0 and *gun1G*/Col-0, and absent in NF/MS or LIN/MS. Concerning editing changes, the *ndhB-830* and *ndhB-872* editing sites were weak candidates, while *rpl23-89* was a stronger candidate. However, the editing efficiency of *rpl23* is reduced under stresses (3), speaking for a pleiotropic effect. Additionally, editing of *rpl23* was not progressively restored in *gun1M* seedlings. Regarding splicing alterations, potential affected loci included *ndhA*, *ndhB*, *rpoC1*, *rps12B*, *clpP_1* (no progressive rescue observed for *clpP_1*), and *rpl2* (weak; see fig. S7).

**(C)** Snapshots across the *rps12B* gene. The read depths were visualized with the Integrated Genome Browser (IGB). Splicing for *rps12B* was still observed, and notably, *rps12B* transcripts were largely decreased.

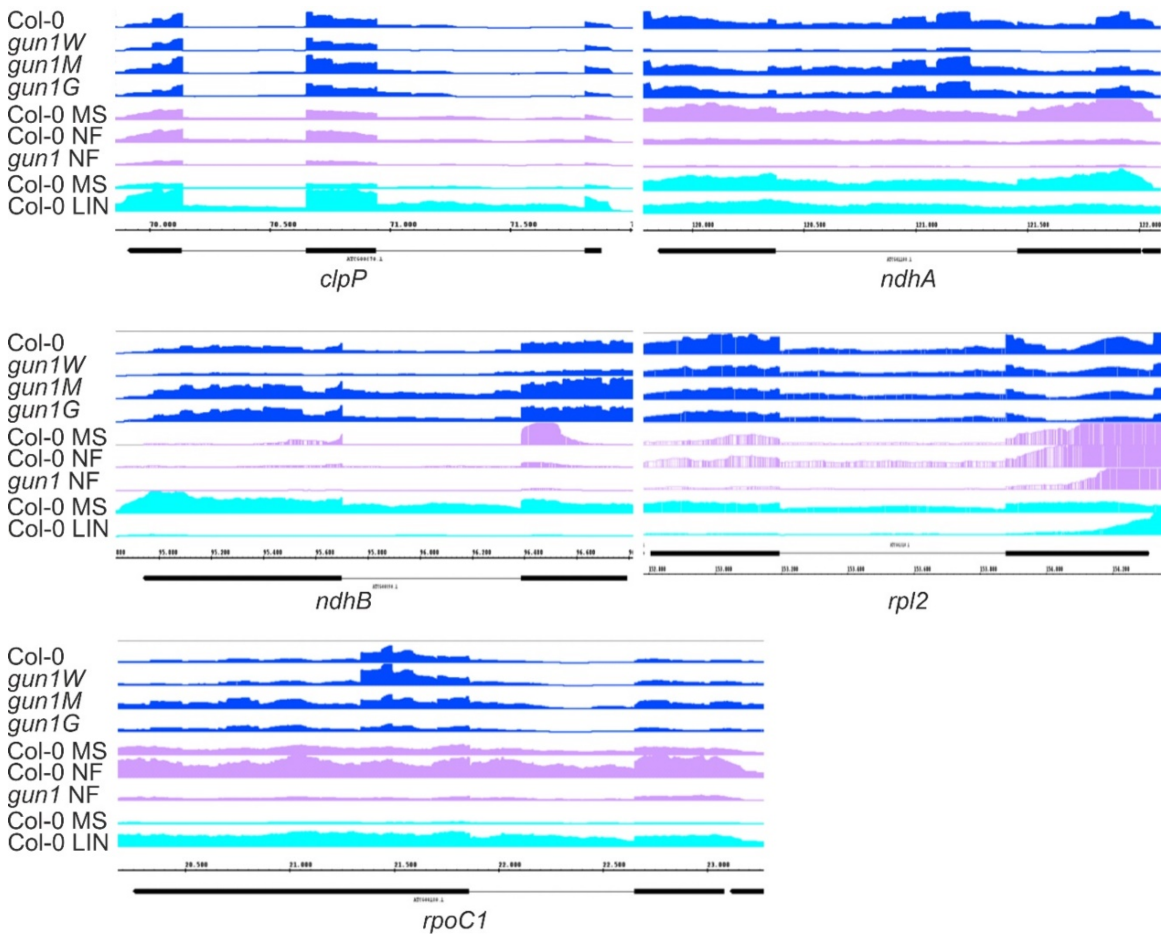


**Fig. S6. Illustration of splicing behavior of selected transcripts.**

Snapshots across the *clpP*, *ndhA*, *ndhB*, rpl2, and *rpoC1* genes are shown. The read depths were visualized with the Integrated Genome Browser (IGB).

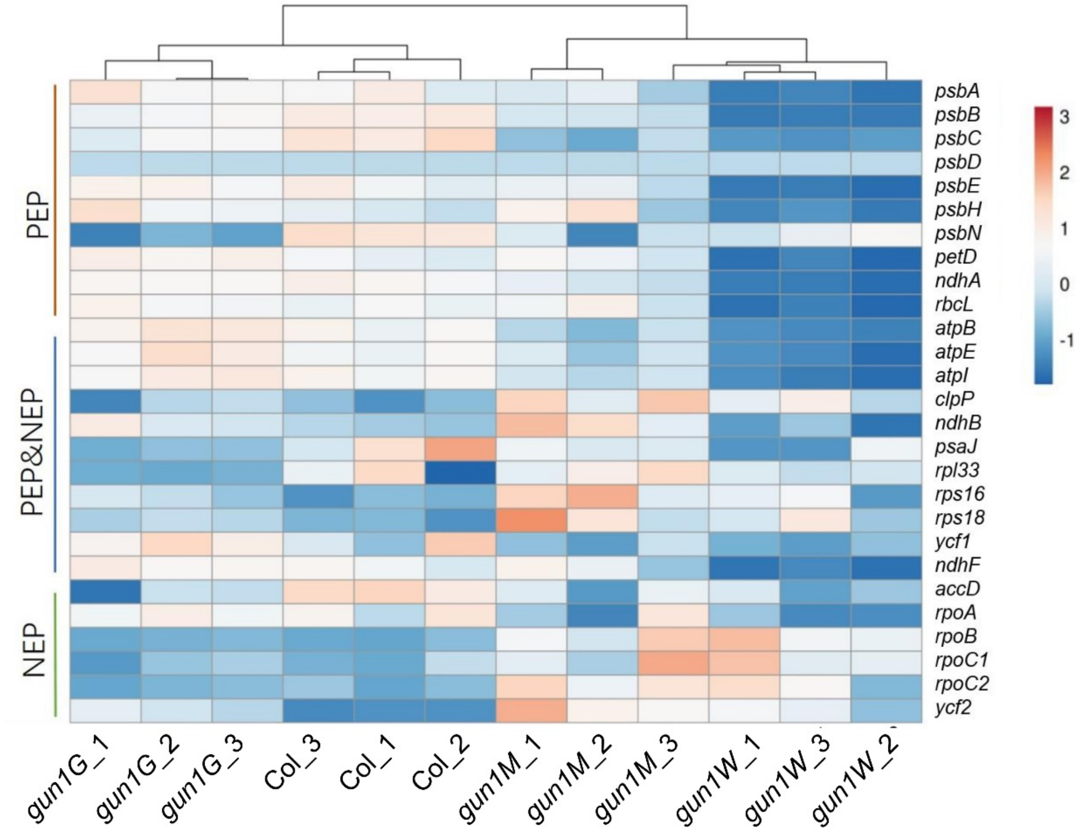


**Fig. S7. Heatmap illustrating the impact of GUN1 deficiency on NEP- and PEP-encoded plastid transcripts (Z-scores).**

Low to high expression is represented by the blue to red transition. Note that Z-scores are calculated for each individual transcript over the different genotypes. NEP is a single-subunit enzyme, whereas PEP consists of core subunits that are encoded by the plastid genes *rpoA*, *rpoB*, *rpoC1* and *rpoC2* (which are transcribed by NEP), and additional protein factors (sigma factors and polymerase-associated proteins, PAPs) encoded by the nuclear genome (4, 5). The general picture was that only PEP transcribes photosystem I and II genes (*psa* and *psb*), most other genes have both NEP and PEP promoters, while NEP alone transcribes a few housekeeping genes (*rpoB*, *accD*, *ycf2*) (6). However, more recent analyses show that the division of labor between NEP and PEP is more complex (4, 7), and no clear conclusion can be drawn about PEP- or NEP-dependent transcription in *gun1W*: the so-called PEP-dependent genes were lower in *gun1W* compared to Col-0, as were the genes transcribed by PEP and NEP, although to a lesser extent. NEP-dependent gene expression was also reduced or in the range of Col-0. NEP, nuclear-encoded RNA polymerase; PEP, plastid-encoded RNA polymerase.

**A**

| | | *gun1W* | *gun1M* | *gun1G* | NF | LIN | Pred. target of GUN1 |
|---|---|---|---|---|---|---|---|
| *ATCG00080* | *psbI* | 0.12 | 0.23 | 0.23 | n.d. | red. | no |
| *ATCG00360* | *ycf3* | 0.41 | 1.26 | 1.22 | 0.54 | 0.94 | no |
| *ATCG00380* | *rps4* | 0.45 | 1.01 | 0.93 | 0.77 | 0.73 | no |
| *ATCG00500* | *accD* | 0.38 | 0.68 | 0.73 | 0.62 | 0.56 | no |
| *ATCG00550* | *psbJ* | 0.10 | 0.81 | 0.48 | n.d. | red. | no |
| *ATCG00560* | *psbL* | 0.08 | 0.90 | 0.90 | n.d. | red. | no |
| *ATCG00570* | *psbF* | 0.07 | 0.43 | 1.18 | n.d. | red. | no |
| *ATCG00740* | *rpoA* | 0.21 | 0.53 | 1.03 | 1.25 | 2.26 | no |
| *ATCG00750* | *rps11* | 0.19 | 0.63 | 1.14 | 1.28 | 2.20 | no |
| *ATCG00770* | *rps8* | 0.18 | 0.50 | 1.05 | 1.23 | 2.35 | no |
| *ATCG00780* | *rpl14* | 0.15 | 0.58 | 1.39 | 0.87 | 1.84 | no |
| *ATCG00790* | *rpl16* | 0.11 | 0.27 | 0.86 | 0.78 | 1.84 | no |
| *ATCG00800* | *rps3* | 0.18 | 0.68 | 1.50 | 1.03 | 1.35 | no |
| *ATCG00810* | *rpl22* | 0.16 | 0.42 | 1.12 | 1.13 | 1.33 | no |
| *ATCG00820* | *rps19* | 0.13 | 0.35 | 0.77 | 1.12 | 1.45 | no |
| *ATCG00900* | *rps7.2* | 0.09 | 0.25 | 0.74 | 6.86 | 0.20 | no |
| *ATCG00905* | *rps12C* | 0.17 | 0.47 | 0.97 | 0.55 | 0.76 | yes |
| *ATCG00920* | *rrn16S* | 0.05 | 0.16 | 0.41 | 1.77 | 0.00 | no |
| *ATCG01000* | *ycf1.1* | 0.34 | 0.65 | 1.11 | 1.14 | 0.64 | no |
| *ATCG01120* | *rps15* | 0.39 | 0.88 | 1.02 | 0.67 | 1.93 | no |
| *ATCG01130* | *tic214* | 0.18 | 0.40 | 1.53 | 0.85 | 2.44 | yes |

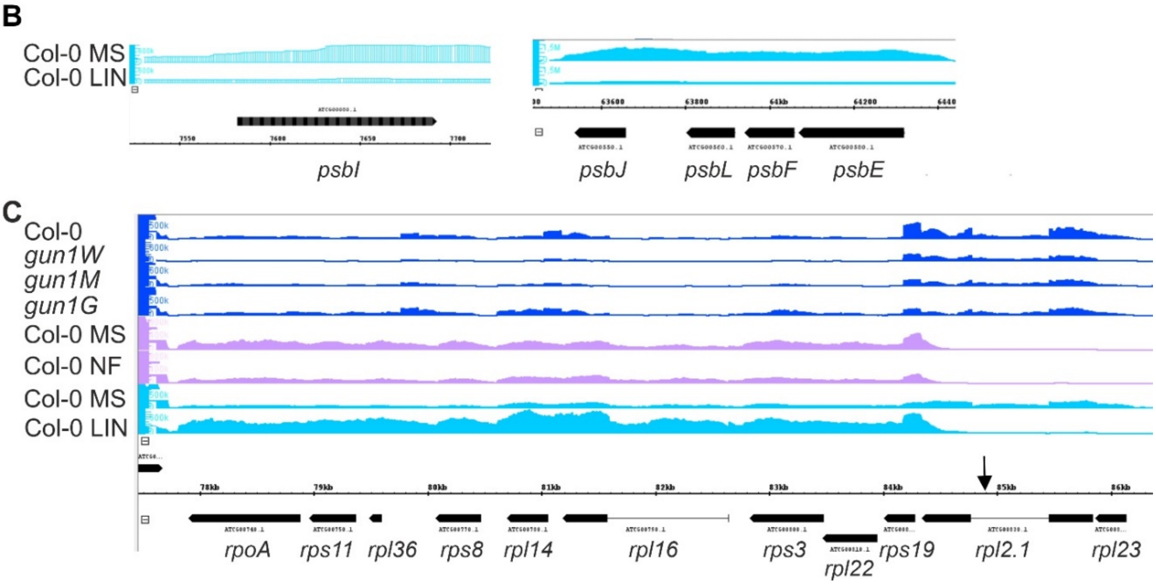


**Fig. S8. GUN1 deficiency has a significant impact on the entire chloroplast transcriptome.**

**(A)** Fold changes of transcripts that are exclusively reduced in *gun1W* are shown. n.d. denotes that these transcripts were not detected in the RNA-Seq analysis; red., reduced.

**(B)** Visualization of reads mapping to the indicated regions in Col-0 grown without (MS) or with lincomycin (LIN).

**(C)** Coverage plots depict the accumulation of reads across the depicted gene cluster.

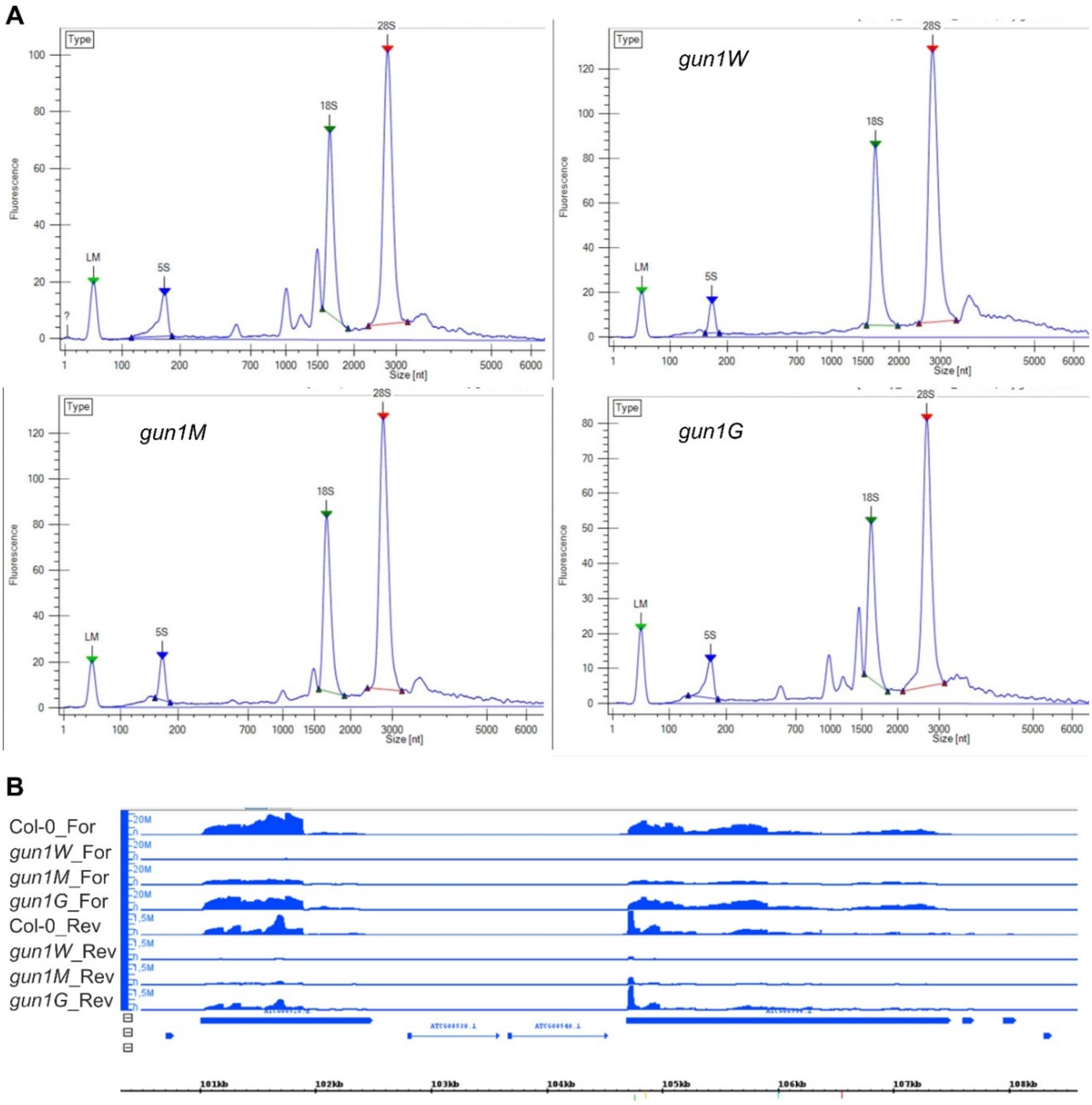


**Fig. S9. Plastid ribosomal RNAs are significantly reduced in *gun1W* and *gun1M* seedlings.**

(**A**) Bioanalyzer profiles of total RNAs prepared with the RNA 6000 Nano Kit (Agilent).

**(B)** Read depths across the ribosomal operon were visualized using the Integrated Genome Browser (IGB). Note that this sequencing technique does not reliably capture tRNAs.

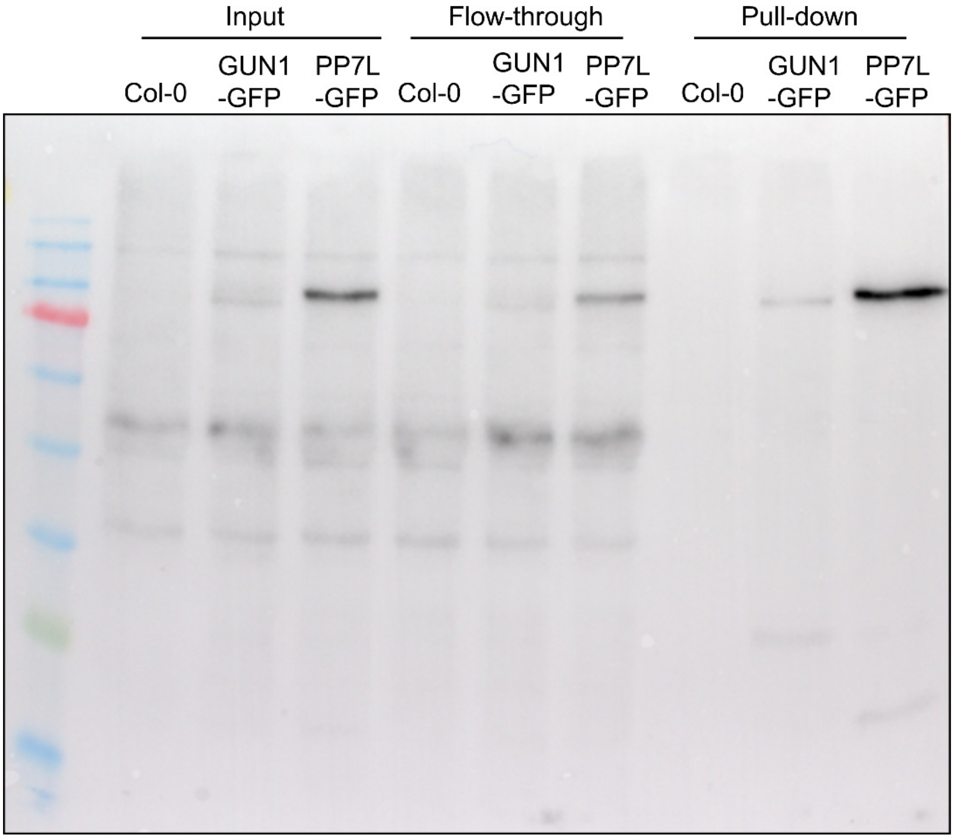


**Fig. S10. Immunoblot analysis for the validation of the GUN1 RIP experiment.**

Immunoblot analysis of proteins isolated from the input, flow-through and pellet (pull-down) fractions of the RIP experiment performed with proteins isolated from 4-day-old Col-0, *35S:GUN1-GFP*, and *35S:PP7L-GFP* seedlings. Proteins were fractionated by SDS-PAGE, and blots were probed with an antibody detecting GFP.

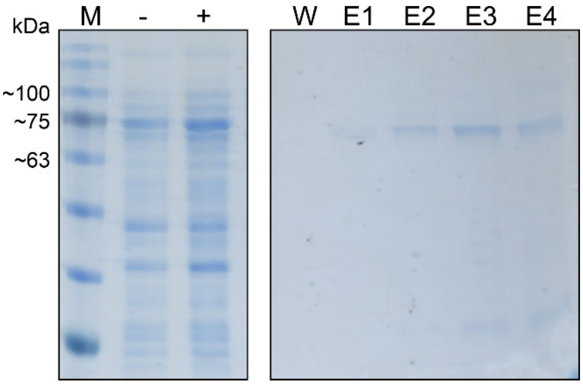


**Fig. S11. Overexpression and purification of a His-tagged GUN1-PS protein in *E. coli*.**

GUN1-PS encompasses all PPR and SMR motifs (PS) spanning amino acids 232 to 918. -, before induction; +, after 20 hours induction at 18°C; W, wash fraction with a buffer containing 20 mM imidazole.; E1 and E2, elution fractions with a buffer containing 250 mM imidazole; E3 and E4, elution fractions with a buffer containing 500 mM imidazole.

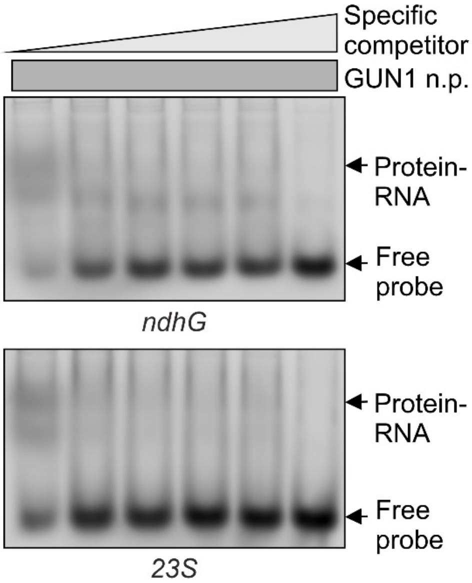


**Fig. S12. The GUN1 protein interacts in vitro with RNA sequences located in *ndhG* and *23S* rRNA.**

EMSAs were performed with His-tagged GUN1-FL protein that was synthesized in vitro. 800 nM of non-purified GUN1 protein (GUN1-FL n.p.) was incubated with 25 nM Cy5-labelled ssRNA probes in the presence of increasing concentrations (indicated by the light-gray triangle) of the same unlabeled ssRNAs as competitor. Binding reactions were then subjected to electrophoresis on nondenaturing TBE-polyacrylamide gels.

**Table S6. Primers used in this study.**

| Atg number | Description | Primer sequence (from 5' to 3') |
|---|---|---|
| RT-qPCR | | |
| AT4G36800 | "Housekeeping" gene RCE1 | CTGTTCACGGAACCCAATTC |
| | | GGAAAAAGGTCTGACCGACA |
| AT2G33430 | MORF2 | ATGGCTTTGCCTTTGTCTG |
| | | AACCTGACCGGTTAGCTC |
| AT1G29910 | LHCB1.2 | CCGTGAGCTAGAAGTTATCC |
| | | GTTTCCCAAGTAATCGAGTCC |
| AT3G01500 | CA1 | GAGAAATACGAAACCAACCCT |
| | | ACATAAGCCCTTTGATCCCA |
| AT1G67100 | PC1 | CAACGCAGGGTTCCCACAT |
| | (primer combination used by Zhao et al. 2019) | CGCACAATAGAAACCGTAAGAGC |
| RNA gel-blot analysis | | |
| AT1G29910 | LHCB1.2 | GACTTTCAGCTGATCCCGAG |
| | | CGGTCCCTTACCAGTGACAA |
| Editing detection by Sanger sequencing | | |
| ATCG00670 | clpP_559 | AATGATCCATCAACCCGC |
| | | ATTGAACCGCTACAAGATC |
| ATCG00300 | psbZ_50 | GGGATTCGAACCCTCGATAG |
| | | TCAAGTTCCATAAGTTCGACCC |
| ATCG00180 | rpoC1_448 (same like Zhao et al. 2019) | TTTTCTTTTGCTAGGCCCATAA |
| | | TTCGCAAATCTAAATCGGCT |
| ATCG00190 | rpoB_338_and_551 | TATCGGTTTATTGATCAGGG |
| | | GCAGCTGCTAACACATCTC |
| ATCG00890 | ndhB_467 | TGCTTCTCTTCGATGGAAG |
| | | TCCTTCGTATACGTCAGG |
| ATCG01010 | ndhF_290 | ACTGCCAGTTATCCAATAAAGAC |
| | | TCATCCCTTTCATTCCACTTC |
| ATCG00065 | rps12_-58 | TGATTAGGTCATTTACCCTG |
| | | AAATACAAGACAGCCAATCC |
| Editing detection by amplicon sequencing | | |
| ATCG00670 | clpP_559 | TCTTGGAAGCGGAAGAATTACT |
| | | TGAACCGCTACAAGATCAAC |
| | psbZ_50 | CCACCAAGAAGACTAATCCAATCC |

| | | |
|---|---|---|
| | | GCTTTCCAATTGGCAGTTTTTG |
| | rpoC1_448 | AGAAGGCCTAGTATACTGCGA |
| | | TAATAATTCGCAAATCTAAATCG |
| | rpoB_551 | GAAAACCAGTAGGAATATGC |
| | | TCCCCACCTACACAAGAAAATTG |
| | ndhB_467 | CCGATGGAGAGAAGAACCTATG |
| | | TATCCAGATAATAGGTAGGAGC |
| | ndhF_290 | AAAACCTTCGCCGCATGTGG |
| | | ATCAGAACCAAAATCCCAACAG |
| RIP-qPCR | | |
| ATCG00950 | 23S_104766 and 23S_104856 | GATACCTAGGCACCCAGAGAC |
| | | CTACTAAGATGTTTCAGTTCGCCA |
| ATCG00950 | 23S_106558 | CGGAAGGTTAAGGAAGTTGG |
| | | GGAATTTCGCTACCTTAGGAC |
| ATCG01080 | ndhG | TCGATACGTCATGGTACGGG |
| | | TGACGAGCCACAGAAATTGC |
| ATCG01130 | ycf1.2 | ATGTACCAATGGAGCCTGGA |
| | | GGATCAAAGCCATTTCATCGT |
| ATCG00280 | psbC (negative control) | ACTTCCCCACCTAGCCACTT |
| | | AGCCCAAAACTGCAGAAGAA |
| ATCG00020 | psbA (negative control) | TTTCCGGTGCCATTATTCCT |
| | | TCATAAGGACCGCCGTTGTA |
| ATCG00360 | ycf3 (negative control) | CGGATGTCGGCTCAATCTGAAGG |
| | | AGGGGTTTCGTTCTAATGCCCGA |
| EMSA | | |
| ATCG00950 | 23S_104856 | aagagacaaccuggcgaacugaaac |
| ATCG01080 | ndhG_118454 | agaaaaaaaucuguugauaaaugaa |
| | U6 | gggccaugcuaaucuucucug |
| Cloning | | |
| AT2G31400 | *GUN1* cDNA cloned into pEU-E01-His-TEV | TATTTTCAGGGC<u>GATATC</u>atggcgtcaacaccgcctcactgg |
| | For in vitro transcription/translation | GTACCCGGGATC<u>CTCGAG</u>ctacaaagaagaggctgtaaagcaaacg |
